# Genetic determinants facilitating the evolution of resistance to carbapenem antibiotics

**DOI:** 10.1101/2021.02.11.430761

**Authors:** Peijun Ma, Lorrie L. He, Alejandro Pironti, Hannah H. Laibinis, Christoph M. Ernst, Abigail L. Manson, Roby P. Bhattacharyya, Ashlee M. Earl, Jonathan Livny, Deborah T. Hung

## Abstract

In this era of rising antibiotic resistance, in contrast to our increasing understanding of mechanisms that cause resistance, our understanding of mechanisms that influence the propensity to evolve resistance remains limited. Here, we identified genetic factors that facilitate the evolution of resistance to carbapenems, the antibiotic of “last resort,” in *Klebsiella pneumoniae*, the major carbapenem resistant species. In clinical isolates, we found that high-level transposon insertional mutagenesis plays an important role in contributing to high-level resistance frequencies in several major and emerging carbapenem-resistant lineages. A broader spectrum of resistance-conferring mutations for select carbapenems such as ertapenem also enables higher resistance frequencies and importantly, creates stepping-stones to achieve high-level resistance to all carbapenems. These mutational mechanisms can contribute to the evolution of resistance, in conjunction with the loss of systems that restrict horizontal resistance gene uptake, such as the CRISPR-Cas system. Given the need for greater antibiotic stewardship, these findings argue that in addition to considering the current efficacy of an antibiotic for a clinical isolate in antibiotic selection, considerations of future efficacy are also important. The genetic background of a clinical isolate and the exact antibiotic identity can and should also be considered as it is a determinant of a strain’s propensity to become resistant. Together, these findings thus provide a molecular framework for understanding acquisition of carbapenem resistance in *K. pneumoniae* with important implications for diagnosing and treating this important class of pathogens.

## Introduction

Antibiotic resistance is one of the most urgent threats to public health. Resistance has emerged to almost all clinically used antibiotics and in nearly all bacterial pathogen species. Numerous studies have focused on identifying and characterizing resistance mechanisms; meanwhile, our understanding of mechanisms that facilitate the evolution of resistance in clinical isolates is less well understood^1^. As such, antibiotic efficacy as reflected in minimum inhibitory concentrations (MICs) remains almost the sole criterion to guide clinical antibiotic choice. However, more sophisticated antibiotic stewardship that considers the frequency of the evolution of resistance in antibiotic selection would help to preserve the existing arsenal of antibiotics. Such stewardship would need to be informed by an increased understanding of the mechanisms that may affect the evolution of resistance including microbial intrinsic factors such as the genetic background of an isolate and extrinsic factors such as the antibiotic choice.

Bacteria acquire antimicrobial resistance through horizontal gene transfer (HGT) or mutation, processes that can be influenced by intrinsic microbial genetic factors, such as phage defense systems and error prone polymerases, respectively^2,3^. While HGT involves the acquisition of new resistance genes, mutation of existing genes can occur by acquisition of single nucleotide polymorphisms, insertions, deletions, recombination, or transposition events. At the same time, microbe extrinsic factors such as the antibiotic identity can also affect the evolution of resistance, as they vary in their ability to induce mutagenesis^4^, have different barriers to resistance^5^, and vary in their spectrum of possible resistance conferring mutations.

The carbapenems, which are the latest generation of β-lactams, are often used to treat infections resistant to almost all antibiotics including extended spectrum β-lactam antibiotics^6,7^. Carbapenem resistance thus typically emerges in bacteria that already carry extended spectrum β-lactamases (ESBLs) and/or other β-lactamases^8–10^. Carbapenem resistance is most often mediated by the production of carbapenemases. In the absence of carbapenemases however, resistance can be achieved through the acquisition of a combination of porin mutations to impede drug entry and/or significant increases in β-lactamase expression^8–11^. Therefore, the evolution of carbapenem resistance often involves complex mechanisms of HGT and mutation acquisition.

The Gram-negative pathogen *Klebsiella pneumoniae* is one of the most prevalent carbapenem resistant Gram-negative species^12,13^. Within this species, carbapenem-resistance occurs predominantly in a few clonal groups (CG), such as CG258, CG15, and CG20^8,13–18^. While clonal spread plays a role in the dissemination of carbapenem resistance^8,15^, the emergence of new highly resistant lineages^19–24^ and the independent acquisition of carbapenem resistance by distinct CG258 strains^14,20,25,26^ suggest that ongoing evolution of carbapenem resistance also plays an important role. These observations suggest that the underlying genetic background of CG258 and other emerging lineages may contribute to a higher propensity for resistance acquisition. Recently, many bioinformatic studies have reported that a major phage defense system, the CRISPR-Cas system, is absent in Sequence Type ST258 and ST11 strains, two major lineages of CG258^27–29^. As one of the earliest lineages causing outbreaks of carbapenem resistance, ST258 *K. pneumoniae* isolates are responsible for the global spread of *K. pneumoniae* carbapenemases (KPC)^8,15^. Therefore, it has been suggested that the lack of CRISPR-Cas systems could be one of the genetic factors contributing to the high rates of carbapenem resistance in this group. However, the more recently emerging lineages, such as ST15 and ST307 do contain such systems and so carbapenem resistance more generally cannot be explained so simply.

Meanwhile, antibiotic identity may also affect the frequency of evolving resistance. Currently, four different carbapenems are available in an intravenous formulation^6^: imipenem, meropenem, ertapenem, and doripenem. In addition, faropenem, a related oral antibiotic in the penem class, is available but only outside of the U.S.^30^. Although the five drugs share similar structures and mechanisms of action, differences in their pharmacokinetics (ertapenem can be administered once a day while the other carbapenems require administration 3-4 times per day), stability against β-lactamase hydrolysis, and penicillin-binding protein target preference^7,31–33^ may influence the evolution of resistance differently. For example, previous studies have shown that compared to other carbapenems, ertapenem is more susceptible to hydrolysis by some β-lactamases and its cell entry is more impeded by the loss of porins^34,35^, raising the possibility that a broader spectrum of mutations on β-lactamase or porin genes may selectively affect ertapenem but not the other carbapenems.

In this study, to understand how bacterial genetic background and different carbapenems affect the rates of resistance evolution, we compared mutation frequencies (previously defined as the frequency of independent resistant mutants emerging in a given population^36^) of carbapenem-susceptible *K. pneumoniae* clinical isolates from ten lineages and found that isolates from the dominant and emerging carbapenem-resistant lineages had higher mutation frequencies leading to carbapenem resistance than other lineages. We demonstrated that the higher mutation frequencies are caused by high-level transposon insertional mutagenesis, a process leading to resistance gene duplication and reversible porin disruption. We also showed experimentally that one of the major phage defense systems, CRISPR-Cas systems, indeed can play a role in restricting resistance gene acquisition when corresponding spacers sequences are present. Furthermore, we found that a broad spectrum of resistance-conferring mutations for selected carbapenems such as ertapenem contributed to increased resistance rates; importantly, these mutations selected from ertapenem exposure could serve as stepping stones to high-level resistance to all carbapenems. Taken together, this work identified multiple factors that facilitate the evolution to carbapenem resistance in *K. pneumoniae* clinical isolates, and demonstrates that the evolution of antibiotic resistance can be a complex process with important implications for antibiotic selection tailored to the genetic background of clinical isolates.

## Results

### The evolution of carbapenem resistance was affected by genetic background of the isolates

We analyzed genomes of 267 previously sequenced *K. pneumoniae* clinical isolates^8^ and selected carbapenem-susceptible isolates from ten lineages (Fig. 1, Supplementary Table 1&2). We chose isolates from (1) the predominant carbapenem-resistant lineage that has caused many outbreaks since 2000 (UCI38 (ST258)), (2) the dominant ESBL-producing lineage that is becoming increasingly carbapenem-resistant (MGH222 (ST15)), (3) newly emerging carbapenem-resistant lineages (UCICRE126 (ST147), MGH66 (ST29), BIDMC41 (ST37), MGH74 (ST76), MGH158 (ST152), UCI64 (ST17)), and (4) lineages that have not caused carbapenem-resistant clonal outbreaks (UCI34 (ST34) and MGH21 (ST111)). We measured mutation frequencies of these isolates under ertapenem (Fig. 2a) or rifampicin treatment (Fig. 2b), using a modified Luria-Delbrück system in which low numbers of bacterial cells were seeded into each well of 384-well plates, thus making the emergence of two independent mutants in the same well extremely unlikely^37^ (Supplementary Fig. 1). This format requires that all resistance occurs through mutation acquisition and not HGT. (We define resistance as at least a 2-fold increase in the MIC for the mutant relative to the MIC against the original susceptible parent strain, and not relative to the clinically defined MIC breakpoints of the antibiotic. Therefore, resistant mutants selected from our experiments do not necessarily have MICs that are greater than the clinical breakpoints.) We found that except for MGH66 (ST29), all isolates showed similar levels of mutation frequencies to rifampicin (Fig. 2b), whereas a wide range of mutation frequencies to ertapenem were observed (Fig. 2a). In particular, some strains had much higher mutation frequencies to ertapenem than to rifampicin. Since resistance to rifampicin is acquired through point mutation resulting from errors during DNA replication^38,39^, these results suggest that other genetic mechanisms help to determine the mutation frequency to ertapenem.

**Fig. 1.**
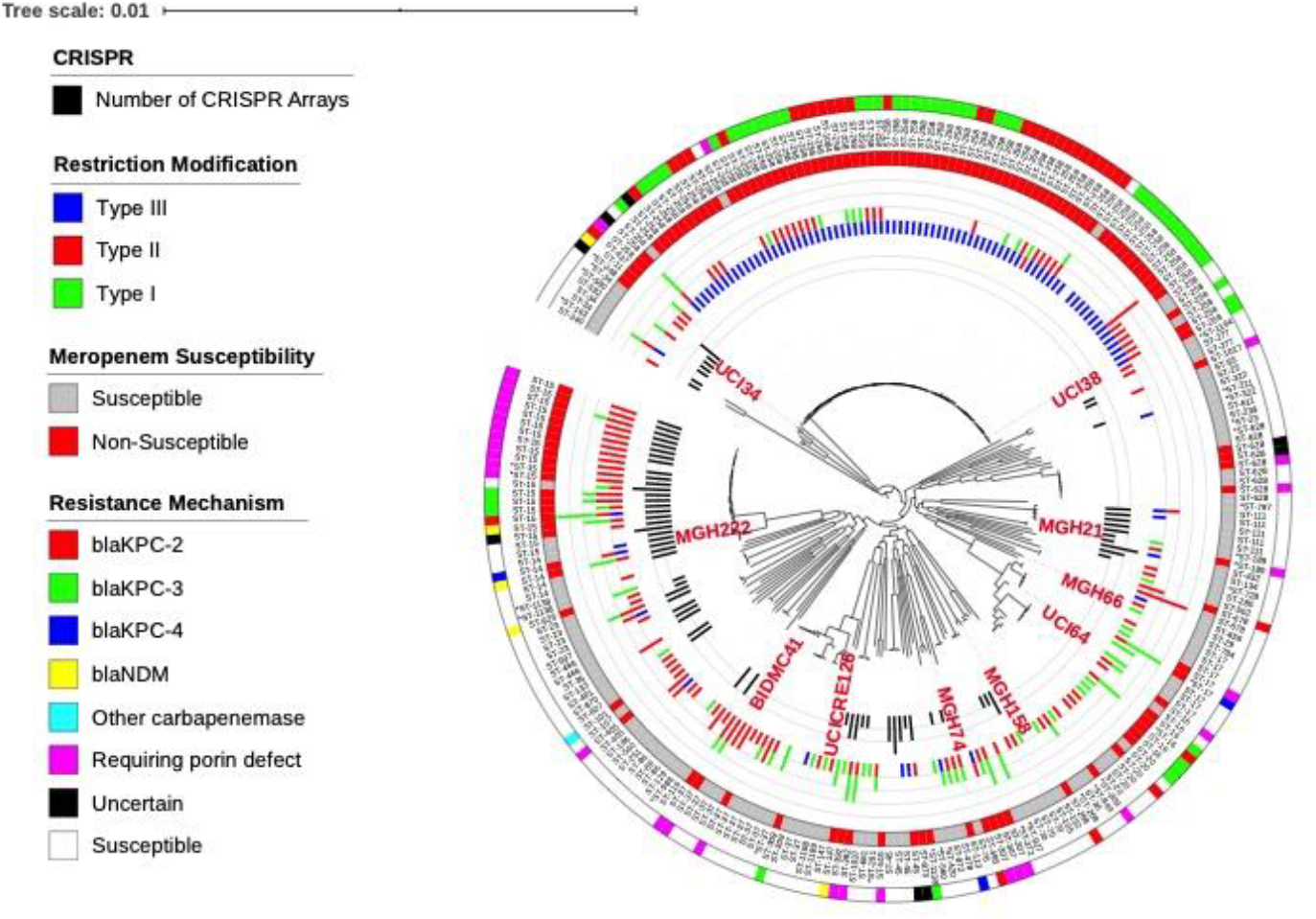
Ten phylogenetically diverse carbapenem-susceptible *K. pneumoniae* isolates were selected from a collection of 267 *K. pneumoniae* clinical isolates. The selected isolates are highlighted in red. In this phylogenetic tree, from inner to outer circles, the content of the CRISPR-Cas systems, Restriction-Modification systems, susceptibility to carbapenems, and sequence types are indicated. For carbapenem-resistant isolates, the resistance mechanism is also indicated.

**Fig. 2.**
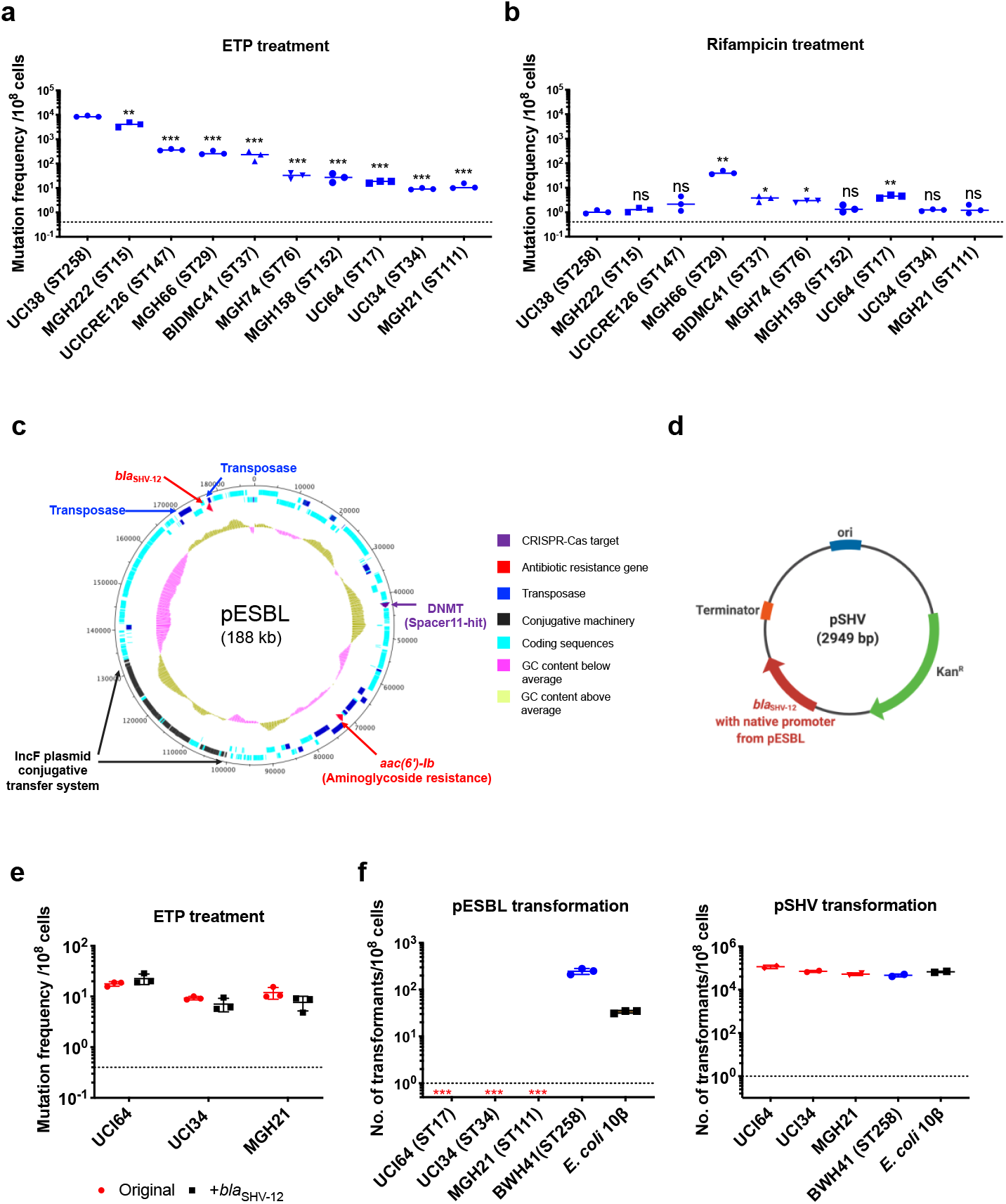
The evolution of carbapenem resistance is affected by genetic background of the isolates. **a**, Mutation frequencies of ten clinical isolates under treatment with ertapenem. Five isolates, UCI38 (ST258), MGH222 (ST15), UCICRE126 (ST147), MGH66 (ST29), and BIDMC41 (ST37) have relatively greater mutation frequencies to ertapenem (>100 mutants per 10^8^ cells) than the other five isolates. Comparing to UCI38 (ST258) which has the highest mutation frequencies to ertapenem, all isolates have significantly different mutation frequencies. Two-tailed Student’s t-test was used for statistical analysis between UCI38 (ST258) and other isolates. **b**, Mutation frequencies of ten clinical isolates under treatment with rifampicin. Isolates with relatively high-level mutation frequencies to ertapenem do not necessarily have high-level mutation frequencies to rifampicin. Two-tailed Student’s t-test was used for statistical analysis between UCI38 (ST258) and other isolates. **c**, Diagram of pESBL, an ESBL-encoding plasmid isolated from UCI38 (ST258). **d**, Diagram of pSHV, a multi-copy laboratory plasmid containing the native promoter and coding region of the ESBL gene *bla*_SHV-12_ amplified from pESBL. **e**, The ESBL gene, *bla*_SHV-12_, was amplified from pESBL and expressed in three isolates lacking an ESBL gene and with relatively low-level mutation frequencies to ertapenem. However, mutation frequencies to ertapenem were not changed compared to the original strains lacking an ESBL gene (red). Two-tailed Student’s t-test was used for statistical analysis to compare the original strain with the corresponding strain overexpressing *bla*_SHV-12_, with p > 0.05 for all three pairs. **f**, Transformation efficiencies of pESBL(left) or pSHV (right) in three isolates lacking ESBL genes (red) and with relatively low-level mutation frequencies to ertapenem. As controls, these two plasmids were also transformed into another ST258 strain BWH41 (blue) which does not carry ESBL genes, and a strain of *E. coli* 10β (black). pESBL could not be transformed into these three isolates but it could be transformed into BWH41 (ST258) and *E. coli*. In contrast, the laboratory construct pSHV was successfully transformed into all strains tested. For all experiments in **a, b, e, f**, two to three independent biological replicates were performed. Data from independent experiments were plotted individually with error bars plotted as the standard deviation. The limit of detection is indicated with a dashed line, and the asterisk (*) under the dashed line indicates frequencies under the limit of detection. * p < 0.05; ** p < 0.005; *** p < 0.0005; ns, not significant.

Among all strains tested, UCI38 (ST258) had the highest mutation frequency to ertapenem. It carries an ESBL gene *bla*_SHV-12_ on the plasmid pESBL (Fig. 2c), raising the possibility that ESBL activity could contribute to high-level mutation frequencies. To test this hypothesis, we transformed pSHV (Fig. 2d), a multi-copy laboratory plasmid containing *bla*_SHV-12_, amplified from pESBL, into three isolates lacking an ESBL gene and with baseline low-level mutation frequencies to ertapenem, including UCI64 (ST17), UCI34 (ST34), and MGH21 (ST111). However, introduction of *bla*_SHV-12_ did not change the mutation frequencies of these strains for ertapenem (Fig. 2e), even though the expression of *bla*_SHV-12_ was higher in strains transformed with pSHV than in UCI38 which naturally carries *bla*_SHV-12_ (Supplementary Fig. 2). This ruled out the simple presence of the ESBL gene alone as the reason for the differing mutation frequencies.

Next, we sought to test the hypothesis that the whole plasmid, pESBL (Fig. 2c), might confer high-level mutation frequencies to ertapenem. However, when we attempted to transform pESBL into the same three strains with low-level mutation frequencies to ertapenem, none of them could take up pESBL. In contrast, an ST258 strain BWH41 (the only ST258 isolate lacking an ESBL gene in our collection) and a laboratory strain of *E. coli*, 10β, could take up pESBL (Fig. 2f). Meanwhile, all strains successfully took up pSHV with similar efficiencies, suggesting that pESBL was uniquely restricted in particular strains.

### A Type I-E CRISPR-Cas system prevented the acquisition of antibiotic resistance genes via HGT while other genetic factors contribute to high mutation frequencies

To understand why pESBL is restricted in these three isolates but not BWH41(ST258), we analyzed the genomic sequences of the collection of 267 *K. pneumoniae* isolates for the presence of two major phage-defense systems, the CRISPR-Cas systems and Restriction-modification systems (Supplementary Table 3), which function to exclude foreign DNA. We found that of the 3 strains which could not take up pESBL, MGH21 (ST111) and UCI34 (ST34) have type I CRISPR-Cas systems, while UCI64 (ST17) has no CRISPR-Cas system but carries Type I R-M systems. In contrast, among 80 strains of the ST258 lineage, we found no CRISPR-Cas systems and most strains carry Type III R-M system (Fig. 1 & Supplementary Table 3). When we broadened our analysis to include the genomic sequences of 2453 *K. pneumoniae* strains available in the NCBI database, including 550 ST258 strains, we found that no ST258 strains contain a CRISPR-cas system (Supplementary table 4), confirming that the lack of CRISPR-Cas system is a genetic feature of the ST258 lineage. This finding is consistent with other bioinformatic studies which have tried to link the absence of CRISPR systems in ST258 strains to carbapenem resistance^27–29^. However, there is no clear association between the absence of CRISPR and the more recently emerging carbapenem-resistant lineages (Fig. 1 and Supplementary Table 3&4).

To understand the ability of MGH21 (ST111) to restrict pESBL uptake, a strain which encodes a Type I-E CRISPR-Cas system but no R-M systems, we first confirmed by RNA sequencing (RNA-seq) that indeed the CRISPR-Cas system was expressed in MGH21 (Supplementary Fig. 3). We then compared the sequence of pESBL with MGH21’s CRISPR-Cas system and found that MGH21 has a spacer (spacer 11) (Fig. 3a & Supplementary Table 5) targeting a gene encoding a DNA-methyltransferase (DNMT)^40^ in pESBL (Fig. 2c); by searching a curated plasmid database^41^, we found that this spacer additionally aligns with sequences found in an additional 94 other plasmids carrying antibiotic resistance genes, including 62 multi-drug resistance plasmids (plasmids carrying resistance genes to more than one class of antibiotics) and 21 plasmids carrying carbapenemase genes (Supplementary Table 6 and 7). In addition, spacer24 (Fig. 3a & Supplementary Table 4) aligned to a conserved hypothetical gene that was also found in UCI38 as well as 66 additional plasmids carrying antibiotic resistance genes, including 44 multi-drug resistant plasmids and 12 plasmids carrying carbapenemase genes (Supplementary Table 6 and 8). Collectively, these results pointed to the potential role of the CRISPR-Cas system in excluding the uptake of resistance carrying plasmids such as pESBL. Indeed, after depleting the CRISPR-Cas operon (MGH21Δ*cas;*the CRISPR-Cas system along with two adjacent hypothetical genes was deleted), pESBL could now be successfully transformed, whereas episomal complementation of the CRISPR-Cas system back into MGH21Δ*cas* again restricted pESBL transformation (Fig. 3b).

**Fig. 3.**
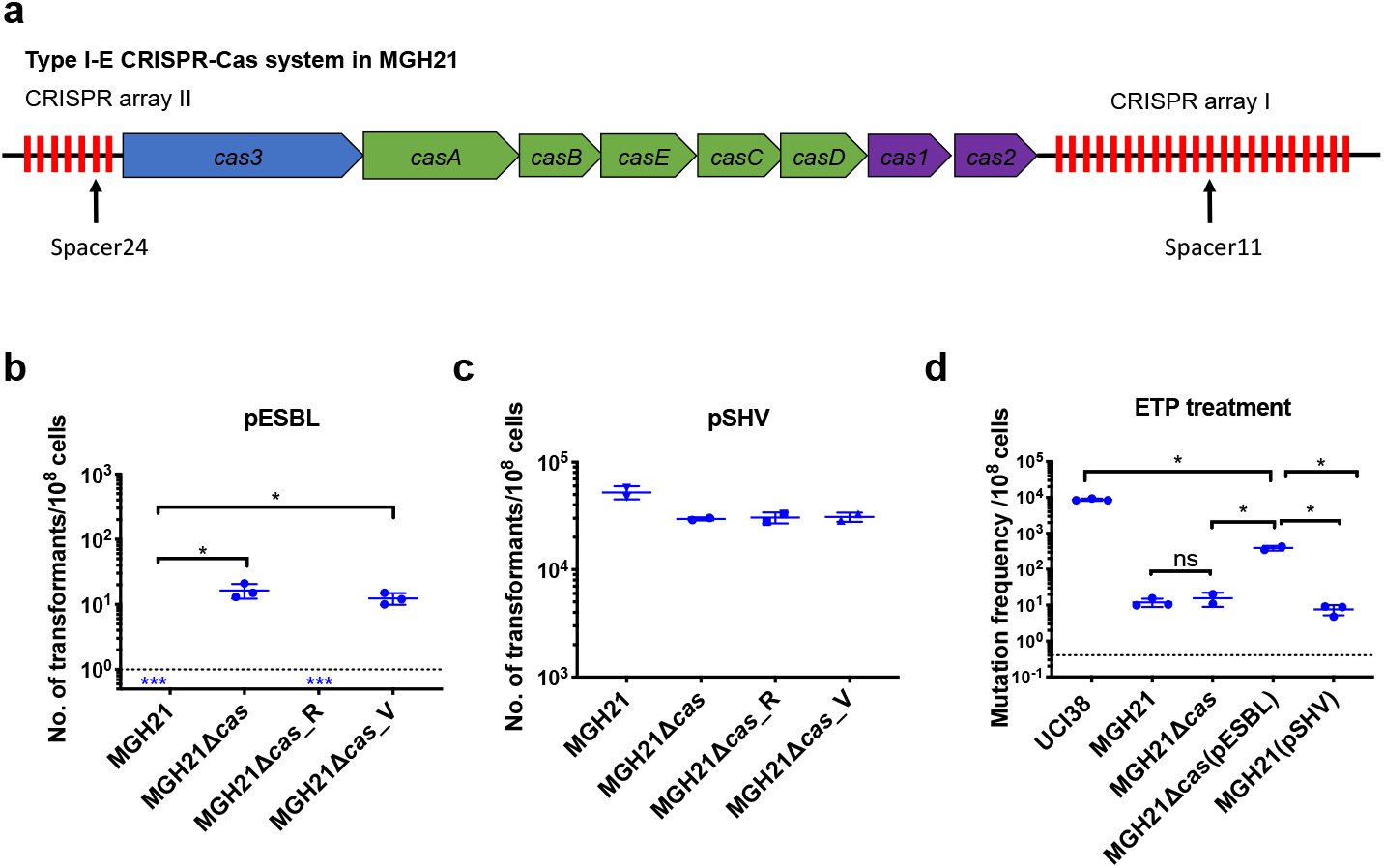
Type I-E CRISPR-Cas system in MGH21 (ST111) prevents the acquisition of pESBL but the presence of pESBL alone does not account for high mutation frequencies. **a**, Type I-E CRISPR-Cas system in MGH21 (ST111). The two CRISPR arrays and the position of two spacers (Spacer11 and Spacer24) that align to plasmids encoding resistance genes are indicated. Spacer11 aligns to the DNMT gene located on pESBL (Fig. 2C). **b-c,** Transformation efficiencies of pESBL (**b**) or the lab construct pSHV (**c**) in MGH21, MGH21Δ*cas*, MGH21Δ*cas*(pCas) with CRISPR-Cas complementation, and MGH21Δ*cas*(pVector) with control vector complementation. pESBL could only be transformed into MGH21 strains in which the CRISPR-Cas system was deleted (MGH21Δ*cas* and MGH21Δ*cas* (pVector)), whereas pSHV could be transformed into all strains at similar efficiencies. **d**, Mutation frequencies of UCI38 (ST258), MGH21, MGH21Δ*cas*, MGH21Δ*cas*(pESBL), and MGH21 (pSHV) with ertapenem treatment. The deletion of the CRISPR-Cas system (MGH21Δ*cas*) and the introduction of pSHV (MGH21(pSHV)) did not affect the mutation frequencies. In contrast, the introduction of pESBL (MGH21Δ*cas*(pESBL)) increased mutation frequencies, indicating that some factors on pESBL other than the ESBL gene affect the mutation frequencies. However, mutations frequencies of MGH21Δ*cas*(pESBL) were still significantly lower than these of UCI38, indicating that more factors in the genetic background of UCI38 contribute to the high-level mutation frequencies. All experiments were performed in triplicate and data were plotted individually. Error bars were plotted as standard deviation. The limit of detection of each assay is indicated with a dashed line, and the asterisk (*) under the dashed line indicates that the transformation efficiencies are below the limit of detection. Two-tailed Student’s t-test was used for all statistical analysis; an asterisk marking a pair-wise comparison denotes a p<0.05.

Unsurprisingly, the absence of CRISPR-Cas system increased rates at which resistance by HGT could be acquired but did not change mutation frequencies of MGH21 (Fig. 3d). In contrast, introduction of pESBL into MGH21Δ*cas* increased the frequency with which resistance to ertapenem emerged in our modified Luria-Delbruck system where HGT cannot occur; the frequency for MGH21Δ*cas* (pESBL) was ~ 30 times higher than for the parent MGH21, MGH21Δ*cas*, or MGH21 carrying pSHV (Fig.3d). As introduction of the ESBL gene alone in pSHV does not change resistance frequencies, this elevation suggests that factors on pESBL other than the ESBL gene contributed to the high mutation frequencies. Further, while MGH21Δ*cas*(pESBL) had elevated ertapenem mutation frequencies relative to MGH21, its frequency was still 10-20 times lower than that of UCI38 itself, from which pESBL was isolated (Fig. 3d), suggesting that differences between the genetic backgrounds of MGH21 and UCI38, irrespective of pESBL, play additional roles in high frequency mutation acquisition.

### Transposon insertional mutagenesis caused frequent and reversible inactivation of porin genes leading to ertapenem resistance

To gain insight into other genetic factors that may cause the different levels of mutation frequencies to ertapenem between UCI38 and MGH21, we analyzed whole genome sequencing (WGS) data of laboratory-derived resistant mutants to identify the specific genetic events leading to ertapenem resistance. We compared six ertapenem resistant mutants derived from UCI38 (ST258), five mutants derived from MGH21 (ST111), and five mutants derived from MGH21Δ*cas*(pESBL) (Fig. 4a), We found that the two strains carrying pESBL favored transposition events as a mechanism to attain resistance while the strain lacking pESBL, MGH21, developed resistance only through SNP acquisition. All six resistant mutants derived from UCI38 were due to duplication of the transposon on pESBL in which the *bla*_SHC-12_ is embedded (Fig. 2c) and/or disruption of *ompK36*, one of the major porin genes of *K. pneumoniae* that facilitates carbapenem cell entry, by insertion sequences (ISs, small transposons that only carry the transposase genes). (Although the other porin OmpK35 also facilitates cell entry for carbapenems, we found no resistant mutants carrying mutations in *ompK35*, probably due to the low expression levels of *ompK35* in the growth condition used^42^ (Supplementary Fig. 4) or pre-existing mutations already disrupting *ompK35^15^* in some strains.) Similarly, for MGH21Δ*cas*(pESBL), four mutants stemmed from the same transposon duplication of *bla*_SHV-12_ on pESBL, while the fifth mutant resulted from the acquisition of a SNP in *ompK36*. In contrast, all resistant mutants derived from MGH21 resulted from the acquisition of SNPs or short deletions/insertions, mostly in porin genes and outer membrane protein genes. pESBL thus increased mutation frequencies relative to pSHV because *bla*_SHV-12_ on pESBL lies within a transposon that can be easily duplicated to elevate ESBL expression and thus MIC (Fig. 3d). In contrast, while carrying pSHV intrinsically conferred a higher baseline MIC because of its higher *bla*_SHV-12_ expression level (Supplementary Fig. 2), it could not duplicate *bla*_SHV-12_ to further evolve increased MIC, thus explaining its unchanged mutation frequencies relative to the parent MGH21 (Fig. 1e).

**Fig. 4.**
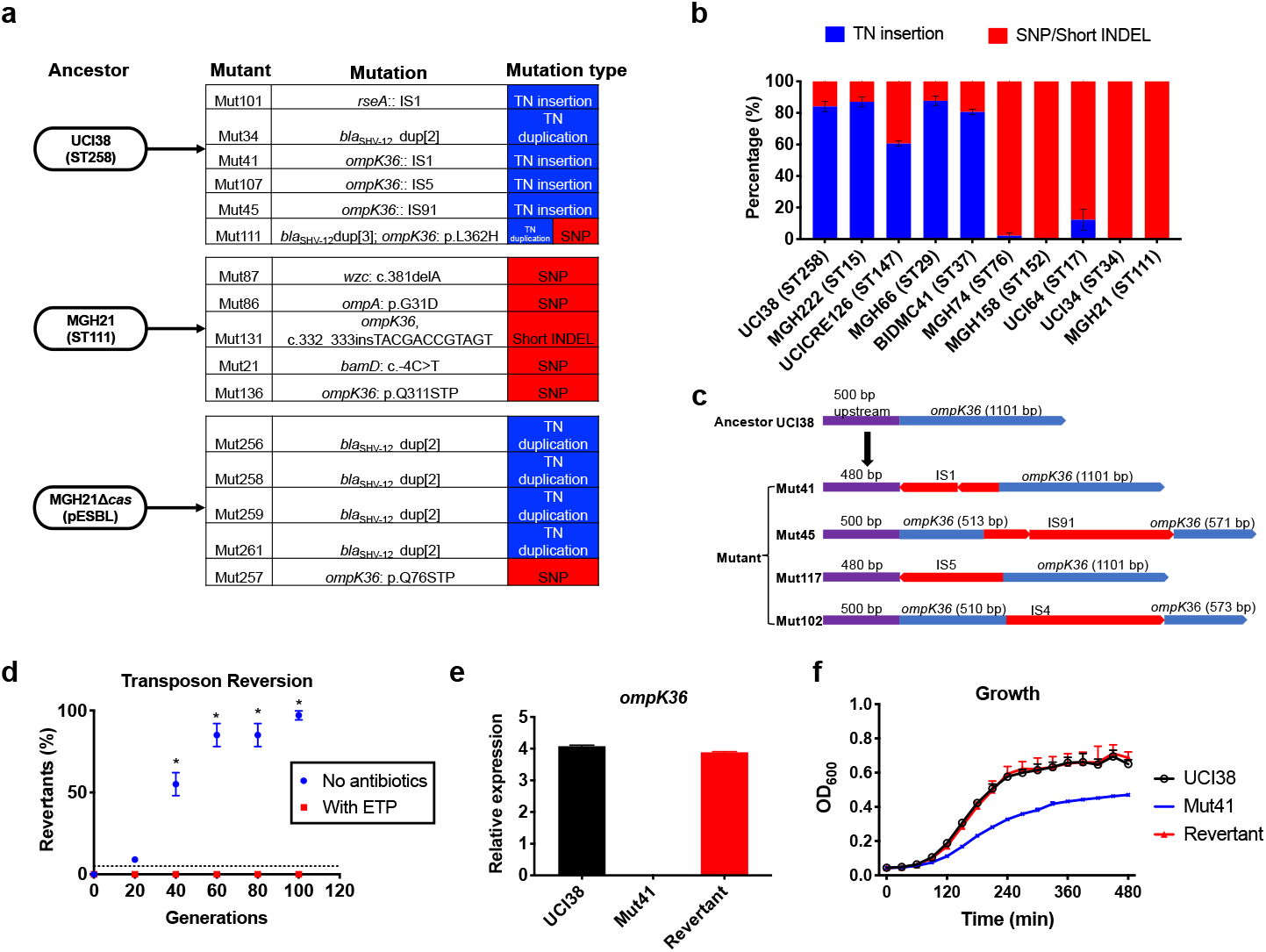
Transposon insertional mutagenesis causes frequent and reversible inactivation of porin genes in isolates with high-level mutation frequencies to ertapenem. **a**, Mutation types (transposon insertion/duplication (blue) vs. SNP (red)) identified via WGS in ertapenem resistant mutants of UCI38 (ST258), MGH21 (ST111), and MGH21Δ*cas*(pESBL). The majority of mutants that carry pESBL (UCI38 and MGH21Δ*cas*(pESBL)) have transposon-mediated mutations, while only SNPs or short insertion/deletions were observed in mutants of MGH21. **b,** Relative quantification of the propensity of ten selected isolates to undergo transposon insertion (blue) versus SNP acquisition (red) in *ompK36* during ertapenem treatment. For each of these strains, 50-100 mutants were isolated, and the types of mutation in *ompK36* locus, if any, was determined via Sanger sequencing. Transposon insertions occurred at ~10 times higher frequencies than the acquisition of SNPs or short insertion/deletion in strains with higher level of mutation frequencies to ertapenem. **c**, Illustration of *ompK36* inactivation by four transposons in UCI38. Four representative mutants derived from UCI38 were selected and the insertion sites were determined by Sanger sequencing. **d,** Transposon disruption of *ompK36* was reversible. A representative ertapenem resistant mutant with an IS1 insertion in *ompK36*, Mut41, was cultured in the presence or absence of ertapenem. Every 20 generations, colony PCR targeting *ompK36* locus was performed on the culture to quantify the percentage of the population that had lost the transposon insertion at this locus. Two-tailed Student’s t-test was used for statistical analysis at each time point to compare the cultures with and without antibiotics. **e-f**, The relative expression of *ompK36* (**e**) and growth curves (**f**) of UCI38 (black), Mut41 (red) and one representative revertant of Mut41 (blue). All experiments were performed in triplicate. Error bars are plotted as the standard deviation.

Comparing the two strains that carry pESBL, we noted that UCI38 was able to disrupt *ompK36* through transposon insertion while MGH21Δ*cas*(pESBL) only did so through SNP acquisition. We hypothesized that the higher likelihood of a disrupting transposition event rather than the acquisition of a disrupting SNP might explain the higher mutation frequencies of UCI38 and other strains with relatively high-level mutation frequencies to ertapenem (Fig. 2a). Indeed, when we used the modified Luria-Delbrück system to isolate and characterize 50 to 100 ertapenem-resistant mutants from each of these ten isolates (Supplementary Fig. 5; Supplementary Table 9), we found that transposon insertions in *ompK36* accounted for 60-90% of resistant mutants derived from strains with high-level mutation frequencies to ertapenem, while only 0-10% of mutants resulted from transposon insertion in *ompK36* in strains with relatively lower mutation frequencies to ertapenem (Fig. 4b). Of note, no one specific IS element accounted for the high transposition rates, as ISs from four different families (IS4, IS5, IS91, IS1) were involved in the inactivation of *ompK36* (Fig.4c and Supplementary Table 10). There was also no correlation between the number of ISs and the activity-level of transposon insertional mutagenesis (Supplementary Fig.6). Nevertheless, these results demonstrate that a higher propensity for transposon insertional mutagenesis in some genetic backgrounds was an important contributor to the more facile evolution of ertapenem resistance in some strains, with such events occurring at nearly ten times higher frequency than SNP acquisition.

In contrast to SNP acquisition for which a reversion is extremely rare, transposon insertions can be reversible^43^. Since porin disruption is known to come at a fitness cost in the absence of antibiotic selective pressure^44,45^, the mechanism of transposon disruption of *ompK36* to achieve antibiotic resistance in UCI38 afforded a potentially facile path, *i.e*., reversion, to recover from this fitness cost when selective pressure is removed. Indeed, this reversion was observed when we cultured Mut41 (Fig. 4c), a mutant of UCI38 carrying an IS1 insertion in the promoter region of *ompK36*, without antibiotics (Fig.4d). 99% of the population reverted to the wild-type *ompK36* gene by ~100 generations, thereby restoring both the expression of *ompK36* and the fitness of the strain relative to the parent mutant Mut41 (Fig. 4e and 4f). We observed the same phenomenon in mutants derived from three other strains (Supplementary Fig. 7) demonstrating the high versatility of this resistance mechanism. A high propensity for transposon insertional mutagenesis resulting in porin inactivation provides a fitness advantage in the presence of antibiotic, while preserving a path to restoration of fitness in the absence of antibiotics.

### Spectrum of genetic mutations conferring resistance to ertapenem is broader than to meropenem

Next, we explored how different carbapenems affect the rates at which resistance evolves. We measured mutation frequencies in response to treatment with four carbapenems and faropenem in three representative carbapenem-susceptible *K. pneumoniae* clinical isolates: UCI38 (an ST258 strain carrying one chromosomal ESBL *bla*_SHV-12_ and a second episomal *bla*_SHV-12_ copy), MGH21 (an ST111 strain with a single copy of the non-ESBL *bla*_SHV-11_ on the chromosome), and MGH32 (an ST111 strain with no β-lactamase genes because the single native, chromosomal *bla*_SHV-1_ is inactivated) (Fig. 5a and Supplementary Table 1). The lowest mutation frequencies resulted from meropenem treatment while relatively higher frequencies resulted from ertapenem and faropenem. In the case of MGH32, which carries no β-lactamase gene, we did not isolate resistant mutants to any of the carbapenems including ertapenem, but isolated resistant mutants to faropenem (Fig. 5a), indicating that β-lactamase genes may be necessary for the evolution of resistance to carbapenems but not to faropenem. To confirm that our observation was not limited to these three strains, we measured mutation frequencies of an additional three isolates under separate treatment of these five antibiotics, and similar patterns were observed, suggesting that the influence of carbapenem identity is independent of the genetic background of strains (Supplementary Fig. 8).

**Fig. 5.**
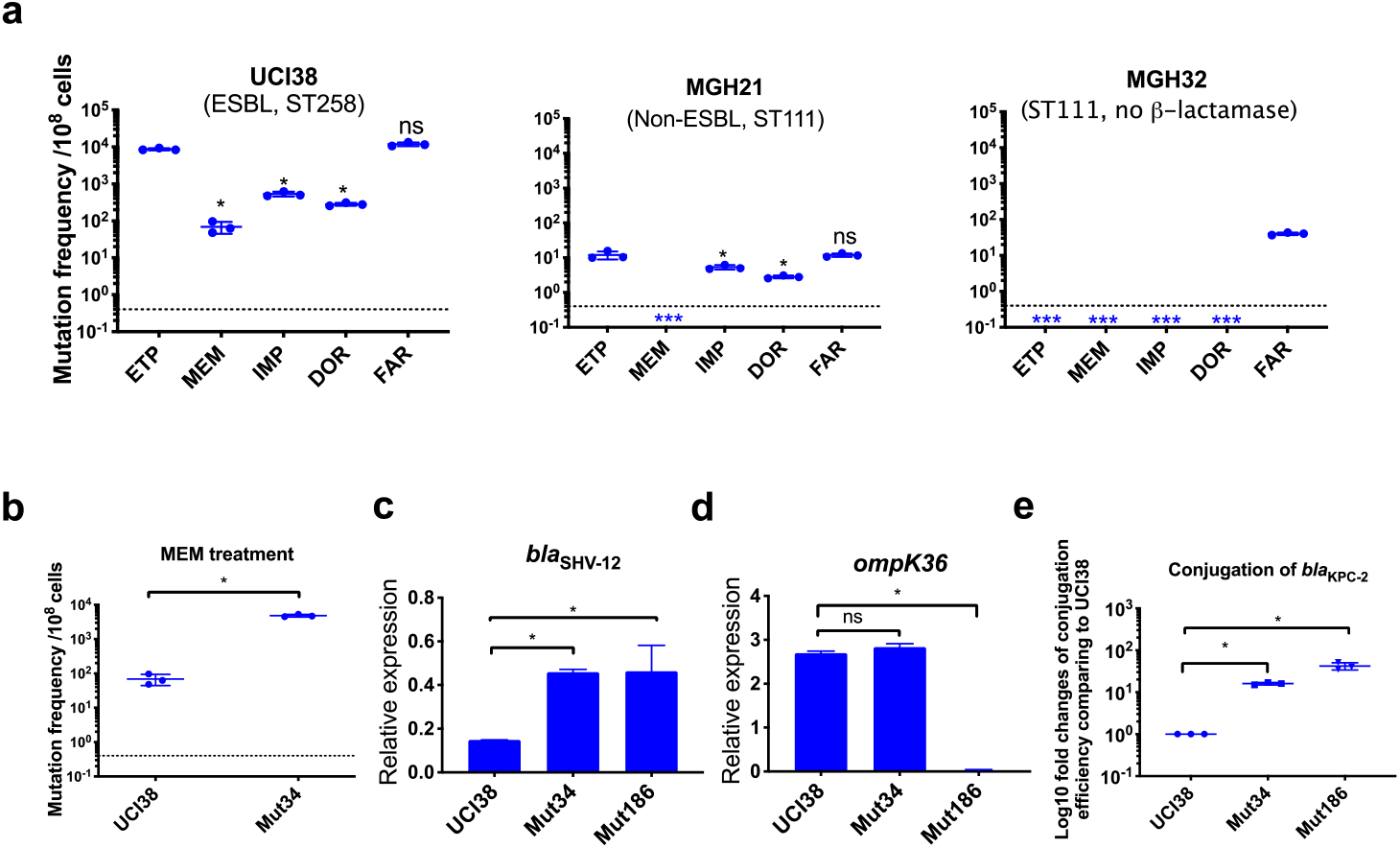
Ertapenem and faropenem treatment are not only associated with higher mutation frequencies, but they also promote the evolution of meropenem resistance. **a**, Mutation frequencies of three representative isolates, UCI38 (ST258), MGH21 (ST111) and MGH32 (ST111, no β-lactamase), under separate treatment with ertapenem (ETP), meropenem (MEM), imipenem (IMP), doripenem (DOR), or faropenem (FAR). Higher mutation frequencies are associated with ertapenem and faropenem treatment, while lower mutation frequencies are observed with meropenem treatment. In MGH32, an isolate without β-lactamase genes, only faropenem-resistant mutants were isolated. Two-tailed student’s t-test was used for statistical analysis to compare between ertapenem treatment and other carbapenems or faropenem. **b**, Mutation frequencies of UCI38 and Mut34, an ertapenem-restricted resistant mutant derived from UCI38, under treatment with meropenem. Despite having the same MIC of meropenem as UCI38, Mut34 had higher mutation frequencies than UCI38. **c-d**, Relative expression levels of *bla*_SHV-12_ (**c**) or *ompK36* (**d**) in UCI38, Mut34, and Mut186 (an ertapenem and meropenem resistant mutant derived from Mut34) show the progressive acquisition of mutations to achieve meropenem resistance. Mut34 has increased *bla*_SHV-12_ relative to its parent UCI38; Mut186 has disrupted *ompK36*, relative to its parent Mut34. **e**, Conjugation efficiencies of UCI38, Mut34, and Mut186 with *K. pneumoniae* clinical isolate BIDMC45 carrying *bla*_KPC-2_. In the presence of meropenem, Mut186 had the highest conjugation efficiency with UCI38 having the lowest. All experiments were performed in triplicate. Two-tailed student’s t-test was used for statistical analysis to compare UCI38 with other strains. Error bars are plotted as standard deviation. The limit of detection is indicated with a dashed line, and the asterisk (*) under the dashed line indicates frequencies under the limit of detection.

Because ertapenem and meropenem were equally stable under these assay conditions (Supplementary Fig. 9a), and bacteria were treated with concentrations of antibiotic normalized to their MICs for each drug, the different mutation frequencies were not explained by differences in antibiotic exposure. We also ruled out the possibility that ertapenem could induce more mutagenesis than meropenem, a phenomenon that has been described for some β-lactams^46^, by measuring the mutation frequencies to rifampin after pre-treatment with sub-MIC concentrations of ertapenem, meropenem, or ciprofloxacin (a fluoroquinolone antibiotic known to induce mutagenesis^4^) as a positive control. While both carbapenems increased rifampin mutation frequencies compared with untreated controls, each did so equivalently, and less than ciprofloxacin (Supplementary Fig. 9b).

We then turned to the possibility that ertapenem’s higher mutation frequency could be due to a greater spectrum of resistance-conferring mutations than for meropenem. We isolated and characterized 90 mutants, derived from UCI38 or MGH21, that were selected from our modified Luria-Delbrück system with confirmed shifts in the corresponding MICs of ertapenem and meropenem (Table 1 and Supplementary Table 10). Sixty-three mutants had increases in the MICs, relative to their corresponding ancestor strains, of both ertapenem (2-256 folds increases) and meropenem, albeit with relatively lower levels of meropenem resistance (2-16 folds increases). We did not isolate any mutants that are highly resistant (MIC > 4μg/mL) to meropenem. Meanwhile, 27 mutants only had corresponding increases in the MICs of ertapenem, and not meropenem (Supplementary Table 11). No mutants had an increased MIC of meropenem but not ertapenem.

We analyzed WGS data from ten representative mutants, five that had MIC shifts to both ertapenem and meropenem, and five that had MIC shifts only to ertapenem (Table1), and validated all identified resistance-conferring mutations by complementation (Supplementary Table 12). Six of the mutants contained either transposon insertions or SNPs in *ompK36* or duplication of *bla*_SHV-12_. Interestingly, four mutants carried novel mutations, including mutations in *wzc* (capsule synthesis), *ompA* (porin), *resA* (anti-sigma E factor), and the promoter region of *bamD* (outer membrane protein assembly factor), with the first three resulting in selective ertapenem resistance. These results show that indeed ertapenem had a wider allowable spectrum of resistance-conferring mutations than meropenem, which yielded a higher mutation frequency.

**Table 1.**
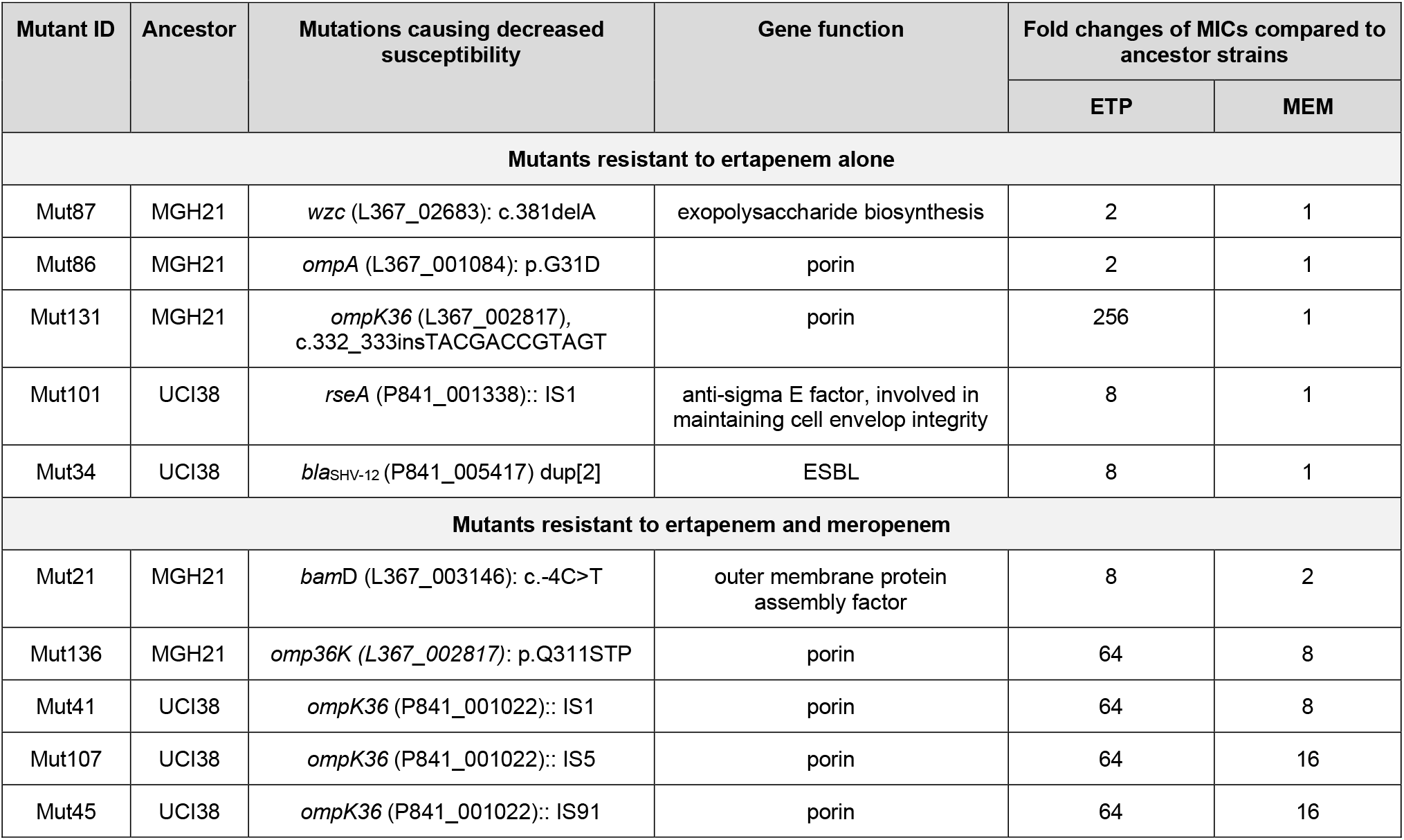
Characterization of representative mutants resistant to both ertapenem and meropenem or to ertapenem alone.

| Mutant ID | Ancestor | Mutations causing decreased susceptibility | Gene function | Fold changes of MICs compared to ancestor strains |  |
| --- | --- | --- | --- | --- | --- |
|  |  |  |  | ETP | MEM |
| Mutants resistant to ertapenem alone |  |  |  |  |  |
| Mut87 | MGH21 | *wzc* (L367_02683): c.381delA | exopolysaccharide biosynthesis | 2 | 1 |
| Mut86 | MGH21 | *ompA* (L367_001084): p.G31D | porin | 2 | 1 |
| Mut131 | MGH21 | *ompK36* (L367_002817), c.332_333insTACGACCGTAGT | porin | 256 | 1 |
| Mut101 | UCI38 | *rseA* (P841_001338):: IS1 | anti-sigma E factor, involved in maintaining cell envelop integrity | 8 | 1 |
| Mut34 | UCI38 | *bla*<sub>SHV-12</sub> (P841_005417) dup[2] | ESBL | 8 | 1 |
| Mutants resistant to ertapenem and meropenem |  |  |  |  |  |
| Mut21 | MGH21 | *bamD* (L367_003146): c.-4C>T | outer membrane protein assembly factor | 8 | 2 |
| Mut136 | MGH21 | *omp36K* (L367_002817): p.Q311STP | porin | 64 | 8 |
| Mut41 | UCI38 | *ompK36* (P841_001022):: IS1 | porin | 64 | 8 |
| Mut107 | UCI38 | *ompK36* (P841_001022):: IS5 | porin | 64 | 16 |
| Mut45 | UCI38 | *ompK36* (P841_001022):: IS91 | porin | 64 | 16 |

### Pre-selection with ertapenem increased the likelihood of evolving resistance to meropenem both by spontaneous mutation and HGT

While many ertapenem resistant mutants do not display resistance to meropenem, we found that acquisition of such mutations, while not impacting the immediate efficacy of meropenem as reflected in the MIC, impacted its future efficacy by increasing the frequency at which resistance to meropenem emerges. The mutation frequencies of an ertapenem-restricted resistant strain (Mut34, which carries a duplication of *bla*_SHV_ on pESBL (Table 1)) were more than 100 times greater than the frequency of its corresponding parental strain UCI38 under identical meropenem treatment (Fig. 5b). WGS of the meropenem resistant mutants revealed that the majority of the mutants derived from Mut34 had acquired new mutations in the porin gene *ompK36* (*i.e*., Mut186, Fig. 5c, d), to accompany the previously acquired *bla*_SHV-12_ duplication. These results demonstrate that the previously acquired mutation in Mut34 that confers ertapenem resistance alone, could serve as a stepping-stone to the subsequent acquisition of a porin disrupting mutation to yield meropenem resistance.

Of note, *de novo* mutation acquisition, even in this stepping-stone fashion, resulted in only low to moderate levels of meropenem resistance (4 - 32 folds increase in MIC from the ancestor strains). With the hypothesis that HGT of carbapenemases or additional ESBL genes may be required to evolve truly high-level meropenem resistance, we examined the impact of the ertapenem-limited resistance mutations on the ability to horizontally acquire resistance genes. Indeed, in the presence of meropenem, higher rates of uptake of a clinical plasmid carrying the carbapenemase gene *bla*_KPC-2_ were observed for both Mut34 and Mut186 than the ertapenem sensitive parental strain UCI38 (Fig. 5e); rather than a direct mechanistic impact, this finding is likely due to longer survival times of these mutants in the presence of meropenem compared to the parental strain affording them a greater opportunity to pick up the plasmid, as the conjugation frequencies are the same in the absence of meropenem (Supplementary Fig. 10). A faropenem-limited resistant mutant, Mut101, like Mut34 for ertapenem, also showed elevated mutation frequencies and conjugation efficiencies in the presence of meropenem compared to its parental strain (Supplementary Table 1 and Supplementary Fig. 11). Together these results suggest that ertapenem and faropenem not only elicit more frequent resistance themselves, but they also select for mutations that can increase the rates at which bacteria acquire high-level meropenem resistance.

## Discussion

In this study, we identified genetic factors that facilitate the evolution of carbapenem resistance in *K. pneumoniae* clinical isolates (Supplementary Fig. 12), one of the most alarming antibiotic-resistant pathogens that have emerged due to our limited arsenal against such organisms. We find that high-level transposon insertional mutagenesis and the mutational spectrum for each carbapenem play important roles in increased mutation frequencies. These mutational mechanisms can work in conjunction with loss of systems that restrict horizontal resistance gene uptake, i.e., the CRISPR-Cas systems, to facilitate the evolution of resistance.

We found that isolates of major and emerging carbapenem-resistant lineages indeed have high-level mutation frequencies to carbapenem antibiotics compared to lineages which have not been linked to carbapenem resistance; this is due to high-level transposon insertional mutagenesis in lineages associated with carbapenem resistance. This highlights the notion that the emergence of predominant resistant lineages did not occur through random events and provide genetic markers that signal isolates with high risk of developing resistance. Importantly, this mechanism of acquiring resistance could serve an evolutionary advantage as the disruption of porins by transposons can revert (Fig. 4d), thereby enabling strains to rapidly adapt to fluctuating environments and optimizing their survival in the presence and absence of antibiotic exposure. The fact that many of the more recently emerging lineages, such as ST15 and ST307, have evolved resistance by a combination of ESBLs and porin truncations may potentially point to the relevance of such mutagenic mechanisms. More generally, transposon-mediated gene duplication has been reported to contribute to heteroresistance in many different bacterial species and antibiotic classes^47,48^. This study thus provides further evidence that mutational events mediated by transposons play a critical role in the evolution of antibiotic resistance in parallel with HGT.

Bioinformatic studies have previously suggested a potential relationship between the absence of CRISPR-Cas systems and carbapenem resistance in the ST258 lineage^27–29^. However, as the more recent resistant lineages to emerge still retain CRISPR-Cas systems, the absence of such systems cannot fully explain the emergence of resistance. Here we demonstrated that they indeed can play a role in restricting the uptake of resistance plasmids, if accompanied by appropriate spacers (Fig. 3). Importantly, bioinformatic analysis of spacer sequences, and not simply the presence or absence of a CRISPR-Cas system alone, is needed to understand the functional role of such systems in resistance gene exclusion in the recently resistant lineages.

The mutational spectrum that confers resistance to each carbapenem also affects evolution frequencies. Currently in practice, several factors affect the choice of a specific carbapenem or faropenem in treating a patient, including its availability, spectrum of activity, dosing schedule, route of administration, and cost. Ertapenem is sometimes favored for the convenience of its once-daily dosing, whereas the other three carbapenems all require 3-4 doses per day. However, ertapenem and faropenem lack activity against *Pseudomonas aeruginosa*, thus limiting their use in some infections^7,49^. Besides these factors, mutation frequencies associated with these antibiotics have not been taken into consideration in antibiotic prescription. In this study, we show that a higher resistance frequency is associated with ertapenem and faropenem due to the broader spectrum of resistance-conferring mutations than is allowed for other carbapenems such as meropenem. Importantly, these mutations can serve as stepping-stones to facilitate the evolution of high-level resistance to all carbapenems. As ertapenem or faropenem are often favored for the convenience of its once-daily dosing or oral bioavailability, respectively, these results highlight the non-equivalence of antibiotics even within the same class of antibiotics with respect to the propensity to evolve resistance. It might suggest that the use of carbapenems with a higher barrier to resistance should be favored to prevent the evolution of carbapenem resistance.

Currently, the choice and administration of an antibiotic is based almost solely on the MIC as an indicator of susceptibility. However, this work shows that treating strains with similar MICs with the same antibiotic could have different outcomes with regards to the emergence of resistance. Isolates with diverse genetic backgrounds can have very different mutational frequencies, despite having the same MIC (Supplementary Table 1). Clearly, some genetic mutations pre-selected from ertapenem or faropenem treatment are not sufficient to change MICs of meropenem, but they can significantly increase the likelihood of evolving resistance to meropenem.

In this current era of rising antibiotic resistance, as significant investment is needed in the discovery of new antibiotics, parallel efforts are needed to guide more judicious use of our current available antibiotics to minimize the emergence of resistance. This work suggests that strategies should not only consider current efficacy, but also consider both the genetic backgrounds of strains and antibiotic choice as they impact the potential for erosion of future efficacy. More generally, this work demonstrates that investigating evolutionary drivers of antibiotic resistance can reveal the root causes of resistance evolution, thereby providing a framework to improve current clinical diagnosis and antibiotic selection.

## Methods

### Modified Luria-Delbrück experiment

The robotic, modified Luria-Delbrück system was adapted from a system that was previously described by Gomez et al.^37^. Exponential growth phase bacterial cultures were diluted to roughly 100 cells per 50 μl (2000 cells/ml) in MHB medium. Then the diluted culture was transferred to three to six 384-well microplates (Falcon, cat#. 353962) using Bravo liquid handling platform (Agilent), and each well of these 384-well plates contained 50 μl of the culture. The plates were sealed using BioExcell Film for Tissue Culture (Worldwide life science, cat. # 41061023) and placed in humidified containers at 37 °C. After incubating for three hours, 10 μl cultures were taken from three randomly selected wells and diluted for plating on LB agar plates to quantify cell numbers. Then antibiotics at specified concentrations were added to the wells using Bravo liquid handling platform at specified concentrations. After adding antibiotics, cultures were incubated in humidified containers at 37 °C overnight. The second day morning, OD600 was read using SpectroMax plate reader (Molecular Device) and mutation frequency was calculated using the following equation: 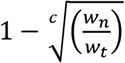, where *c* is the number of cells per well at the time of adding antibiotics, *w_n_* is the number of negative wells, and *w_t_* is the total number of wells. Mutants from each plate were sub-cultured in MHB supplemented with the same antibiotics at the same concentrations used for the selection, and saved in 25% glycerol stocks for future analysis. Mutants that did not grow up in the sub-culturing were excluded from the calculation of mutation frequencies. Each experiment was repeated three times.

To measure mutation frequencies with rifampin after carbapenem treatment, exponential growth phase cultures of UCI38 (OD_600_ ~ 0.2) were diluted 100 times with MHB medium, then the diluted cultures were split into four identical cultures. Cell numbers were quantified by plating diluted cultures on LB agar plates, and these cell numbers were used to calculated mutation frequencies. Ertapenem, meropenem and ciprofloxacin were then added to three of these four cultures at 0.1x MICs. The fourth culture was untreated. Immediately after adding antibiotics, cultures from each condition were aliquoted into three 384-deep-well plates (VWR, cat. # 82051-326) with 50 μl per well using Bravo liquid handling platform. Then these 12 384-deep-well plates were incubated at 37°C with shaking for 2 hours. After incubating for two hours, 10 μl cultures from each well of these 12 deep-well plates were correspondingly transferred to wells of 12 384-clear-bottom microplates (Falcon, cat#. 353962), using Bravo liquid handling platform. Then 50 μl MHB supplemented with rifampin at the concentration of 60 μg/ml was added to each of these wells. The final rifampin concentration in each well was 50 μg/ml. These plates were incubated in humidified containers at 37°C overnight. The second day morning, OD_600_ was read using SpectroMax plate reader (Molecular Device) and mutation frequency was calculated using this equation: 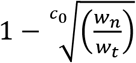, where *c_0_* is the number of cells per well before pre-treatment, *w_n_* is the number of negative wells, and *w_t_* is the total number of wells. This experiment was repeated three times.

### Bacterial strains and plasmid construction

Bacterial strains used in this study are listed in Supplementary Table 1. All strains were cultured in Luria-Bertani (LB) medium or Mueller-Hinton Broth (MHB) with shaking at 37°C or 30°C as specified.

To construct the plasmid pSHV, *bla*_SHV-12_, including the 500 base pairs (bp) upstream region, were PCR amplified from UCI38, respectively, using primers listed in Supplementary Table 13.

Then the PCR products were ligated into vector pSmart LC Kn (Lucigen, cat.# 40821) and electroporated into *E. coli* competent cells 10β (NEB, Cat.# C3020K). Plasmids were then extracted from positive clones and electroporated into *K. pneumoniae* cells that have been made electroporation competent according to the protocol described previously^50^. In brief, *K. pneumoniae* cells were streaked on LB agar plates and grown overnight at 37°C. Then cells were collected directly from LB plates and re-suspended in ice-cold sterilized H_2_O, followed by washing with ice-cold sterilized H_2_O three times. Finally, cells were re-suspended at the concentration of roughly 10^9^ cells/ml for electroporation. Strains expressing *bla*_SHV-12_ were cultivated in medium supplemented with kanamycin at the concentration of 25 μg/ml.

To generate MGH21Δ*cas*, about 1000 bp upstream and downstream of the *cas* operon was amplified from MGH21 using Q5 DNA polymerase (NEB, Cat# M0492). Overlap extension PCR was used to fuse these two pieces of DNA to generate a ~ 2000 bp fragment which was then ligated to pKOV vector^51^ using BamHI and NotI sites, resulting in the construct pKOV-*cas*KO. The construct was transformed to *E. coli* competent cells 10β (NEB, Cat.# C3020K) via electroporation, and the positive transformants were cultured in LB medium supplemented with chloramphenicol (34 μg/ml) at 30°C. Plasmids were then extracted and electroporated into MGH21 electro competent cells and incubated at 30°C on LB agar plates supplemented with chloramphenicol (34 μg/ml) overnight. The integration of the plasmid in either the upstream or the downstream region of the *cas* operon was selected by chloramphenicol resistance and screened by PCR. Following the selection, the integrants were grown in non-selective LB medium for several generations and then plated on LB agar medium with 10% sucrose to induce double recombination. Among the survivors of the sucrose-LB medium, the double recombinants were selected by PCR screening. The deletion of the *cas* operon was confirmed by sequencing and RT-qPCR.

To restore the CRISPR-Cas system to MGH21Δ*cas*, *cas3*, including upstream 500 bp and the CRISPR array II, was amplified from MGH21 and ligated into pSmart LC Kn (Lucigen, cat.# 40821), generating pCas3CRISPR2. Meanwhile, the coding region of *casABECD, cas1, cas2*, and CRISPR array I, were amplified from MGH21 and ligated into pBAD33Gm^52^ using KpnI and XbaI cloning sites, resulting in pBAD33Gm_CasCRISPR1. A SD sequence was also added 8 bp upstream of ATG codon of *casA*. These two constructs were separately transformed into *E. coli* 10β (NEB, Cat.# C3020K) via electroporation. Plasmids were extracted, mixed at 1:1 ratio, and transformed into MGH21Δ*cas*, generating the strain MGH21Δ*cas*(pCas). The transformants containing these two constructs were confirmed using PCR and Sanger sequencing. Similarly, the vector control strain MGH21Δ*cas*(pVector) was generated through co-transforming two empty vectors, pSmart LC KN and pBAD33Gm, into MGH21Δ*cas* strain. When mutation frequencies of MGH21Δ*cas*(pCas) and MGH21Δ*cas*(pVector) with ertapenem were measured in MHB medium supplemented with 1% arabinose (to induce the expression of *casABECD*), kanamycin (25 μg/ml) and gentamicin (10 μg/ml).

### Plasmids extraction and sequencing from UCI38

Plasmids from UCI38 were extracted using QIAfilter Plasmid Midi Kit (Qiagen, Cat.# 12243). Extracted plasmids were then transformed into other clinical isolates and MGH21Δ*cas* through electroporation. Transformants were selected on LB agar plates supplemented with cefotaxime at the concentration of 10 μg/ml. The extracted plasmid DNA was sequenced, assembled and annotated as described before^8^.

### Analysis of 267 *K. pneumoniae* genomes

We used a total of 267 *K. pneumoniae* assemblies generated at the Broad for this analysis, including 80 ST258 strains. *K. pneumoniae* isolates were sequenced, assembled, and annotated as described before^8^. To improve resistance gene predictions, the original gene calls from each assembly were searched against the following databases using BLAST^53^: i) Resfinder^54^ (downloaded January 23, 2018); ii) the National Database of Antibiotic Resistant Organisms (https://www.ncbi.nlm.nih.gov/pathogens/antimicrobial-resistance/; downloaded January 22, 2018); and iii) an in-house database of carbapenemases and ESBLs^8^. For each gene, the database hit with the highest bit score having an e-value <10^-10^ and gene length coverage >=80% was retained. The numbers of annotated carbapenemases and ß-lactamases, including extended-spectrum and broad-spectrum ß-lactamases, were quantified and tabulated for each strain.

### Annotation of Restriction Modification Systems

We downloaded a total of seven reference gene sets for type I (n=3), type II (n=2), and type III (n=2) restriction-modification systems from REBASE (http://rebase.neb.com/rebase/rebase.seqs.html) on May 22, 2019. We used blastn to search for these reference genes in all 267 *K. pneumoniae* assemblies, using an e-value cutoff of 10^-10^ and requiring 80% coverage of the reference gene. We retained the top blast hit for each reference gene set and strain. We considered a restriction modification system of a certain type to be present in a given strain if at least one gene from each of the two (for types II and III) or three (for type I) reference sets were present in the strain.

### Annotation of CRISPR Arrays and *cas* genes

CRISPR Detect^55^ version 2.2 was used to detect CRISPR arrays in the 267 *K. pneumoniae* assemblies and 2453 *K. pneumoniae* strains available in the NCBI database using default parameters. *Cas* genes were identified using the Broad Institute’s microbial annotation pipeline. For the CRISPR arrays identified in MGH21, spacer sequences were aligned to a curated database of plasmid sequences^41^ containing sequences of 6642 plasmids, using blastn and requiring with >80% identity and coverage. Then the sequences of plasmids containing the spacer-hit genes were extracted. ResFinder^54^ was used to identify antibiotic resistance genes in these plasmids, if any, requiring > 95% identity and 80% coverage.

### Determination of MICs

MICs were determined by the broth microdilution method as described^56^. The MICs were measured in duplicates in MHB medium, with a final inoculum size of 5 × 10^5^ cells/ml.

### Quantification of transposon insertions and SNPs in *ompK36*

Following the robotic, modified Luria-Delbrück experiment with ertapenem treatment, 50-100 resistant mutants from each strain were isolated and streaked on LB agar plates supplemented with ertapenem at the concentration of 1.1 x MIC against the ancestor strain. Colony PCR was performed using primers listed in Supplementary Table 13 to amplify *ompK36* locus including upstream 500 bp region of each mutant. The PCR products were then purified and Sanger sequenced. Sequences were aligned to the genomic sequences of the ancestor strains and single-nucleotide variants (SNVs) and transposon insertions could thus be quantified.

### WGS and variant calling

Genomic DNA was isolated using DNeasy Blood & Tissue Kits (Qiagen, cat.# 69504) and quantified using Qubit dsDNA HS Assay Kit (Invitrogen, cat. # Q32851). WGS libraries were made using Nextera XT DNA library preparation kit (Illumina, cat.# FC-131-1096). Then the samples were sequenced using the MiSeq or NextSeq system with 300 cycles, pair-ended. For each strain sequencing depth was set at approximately 100x coverage. BWA mem version 0.7.12^57^, and Pilon v1.23, using default settings^58^, were used to align reads against a reference genome assembly and to identify variants, respectively. SNP positions having mapping quality less than 10 (MQ < 10) were not considered. The *Klebsiella pneumoniae* MGH21, and UCI38 genome assemblies were used as references for variant identification for mutants derived from each respective strain.

### RNA extraction and RT-qPCR

Cells were cultivated in MHB or LB medium at 37°C until early-exponential growth phase. RNA was purified using Direct-zol RNA Kits (Zymo research, cat.# R2070) and quantified with Nanodrop spectrophotometer (ThermoFisher). RT-qPCR was performed using iTaq Universal One-Step RT-qPCR Kits (Bio-Rad, cat.# 1725150). RT-qPCR primers were designed using Primer3^59^ and are listed in Supplementary Table 13. The results were normalized as the percentages of 16 rRNA.

### Reversion of transposon-insertion mutants and growth curves

To check the reverting events of transposon insertion mutants, Mut41, Mut_UCI22, Mut_UCI43, and Mut_UCI44 were cultured in replicates in LB medium with or without ertapenem (1 μg/ml) were set up and diluted every day. Each day, an aliquot of culture (10μL) from each strain/replicate were diluted and plated on LB agar plates to quantify cell numbers. Colony PCR was performed in 24 randomly selected colonies for PCR amplification of the *ompK36* locus, including 500 bp upstream and 100 bp downstream regions. The PCR product was run in agarose gels to assess the size and subsequently Sanger sequenced. One revertant from Mut41 was used for subsequent growth experiment and RT-qPCR to measure the expression of *ompK36*. Growth of UCI38, Mut41 and Mut41_revertant were monitored in a Tecan plate reader in LB medium at 37°C for 8 hours. All experiments were repeated three times.

### Conjugation

Rifampin mutants of UCI38, Mut34, Mut101, Mut186 and Mut195 were raised by plating the exponential-growth phase cells on LB agar plates containing 50 μg/mL rifampin. After overnight incubation, rifampin mutants from each strain were selected and subjected to WGS. Mutants that only have mutations in *rpoB* were selected for conjugation. Exponential growth phase cells of rifampin mutants from these five strains were mixed with BIDMC45 cells at 1:1 ratio, then the mixture was spotted on LB agar plates without antibiotics or containing meropenem (0.003 μg/ml) and grown overnight. The second day morning, cells were transferred to LB liquid medium, serial diluted, and plated on LB agar plates containing meropenem (2 μg/ml) and rifampin (50 μg/ml) for the selection of conjugants. Meanwhile, diluted cells were plated on rifampin (50 μg/ml) plates to quantify cell concentrations. All experiments were repeated three time.

## Data Availability

All data generated or analyzed during this study are included in this article and in the supplementary tables. Sequencing data is deposited to NCBI under the accession number PRJNA670748.

## Acknowledgments

We thank James Gomez for instructions and suggestions of setting up the modified Luria-Delbrück experiment and his further input and comments on this project. We thank Noam Shoresh for the valuable discussions on the data analysis of the modified Luria-Delbrück. We thank Sharon Wong, Anne Clatworthy and Thulasi Warrier for their comments on the manuscript. This publication was supported in part by the National Institute of Allergy and Infectious Diseases of the National Institutes of Health under award 5R01AI117043-05 to DTH and U19AI110818 to the Broad Institute, and by a generous gift from Anita and Josh Bekenstein.

## Author Contributions

P.M. and D.T.H. designed the study and wrote the manuscript. P.M., L.L.H., and H.H.L. performed the experiments. C.M.E. provided suggestions on the initial isolate selection for these experiments and provided critical input on the manuscript preparation. R.P.B. provided extensive input and suggestions on the design of this study and manuscript preparation. A.P. did genomic analysis in the assemblies of 267 *K. pneumoniae* isolates and 2453 *K. pneumoniae* genomes available in the NCBI database. A.L.M. analyzed the WGS data of mutants and identified mutations in each mutant. A.M.E. supervised the genomic data analysis and SNP identification, and provided extensive input on the design of the genomic analysis. J.L. analyzed RNA-seq data. All authors have read and approved the manuscript.

---

**Supplementary Fig. 1.**
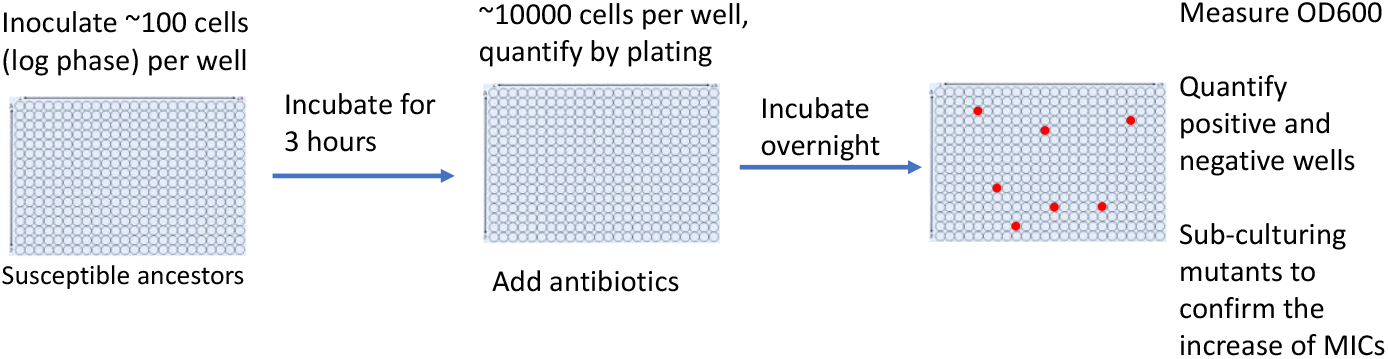
Scheme of the modified Luria-Delbrück system. Exponential-phase growing cells are diluted to roughly 100 cells / 50 μl with MHB medium and inoculated into three to six 384-well plates using Bravo automatic liquid handling platform, followed by incubation at 37°C for 3 hours. 10 μl of culture from 3 randomly selected wells was taken to plate on LB agar plates for estimating numbers of cells in the wells. Then, antibiotics were added at the concentrations of 1.1 x MICs or at specified concentrations using Bravo automatic liquid handling platform, and cultures were incubated at 37°C overnight. OD_600_ was measured the next day and positive and negative wells were quantified. Mutants from each plate were subcultured in MHB medium supplemented with the same antibiotics at the same concentrations used for the selection, and saved in 25% glycerol stocks for future analysis. Mutants that did not grow up in the sub-culturing were excluded from the calculation of mutation frequencies.

**Supplementary Fig. 2.**
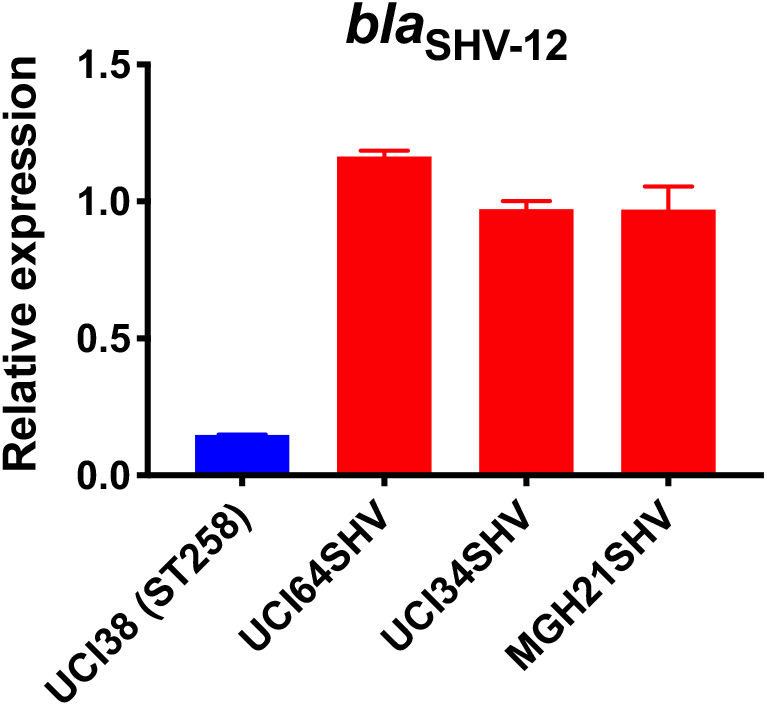
Relative expression of *bla*_SHV-12_ in three strains overexpressing *bla*_SHV-12_ through pSHV. *bla*_SHV-12_, including the promoter region, was amplified from UCI38 and expressed in three strains lacking an ESBL gene. The express levels of *bla*_SHV-12_ in these overexpression strains were higher than it in UCI38 because pSHV is a multi-copy plasmid. RT-qPCR data was normalized to 16S rRNA. Experiments were repeated 3 times and error bars were plotted as standard deviation.

**Supplementary Fig. 3.**
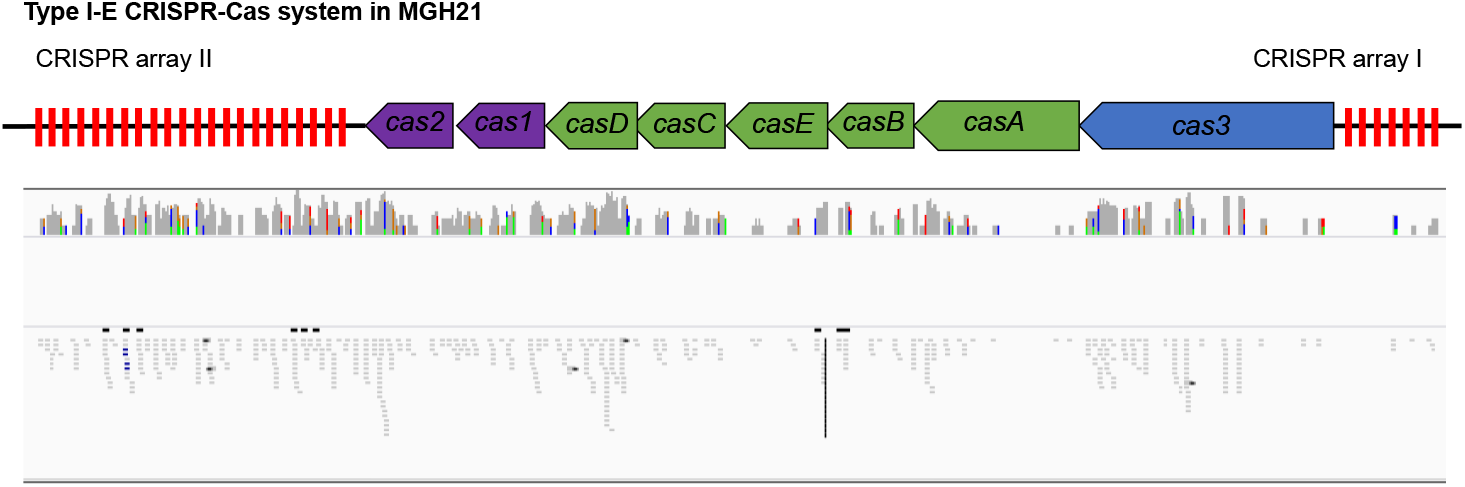
RNA-seq data shows that *cas* genes and most spacers are expressed in MGH21.

**Supplementary Fig.4.**
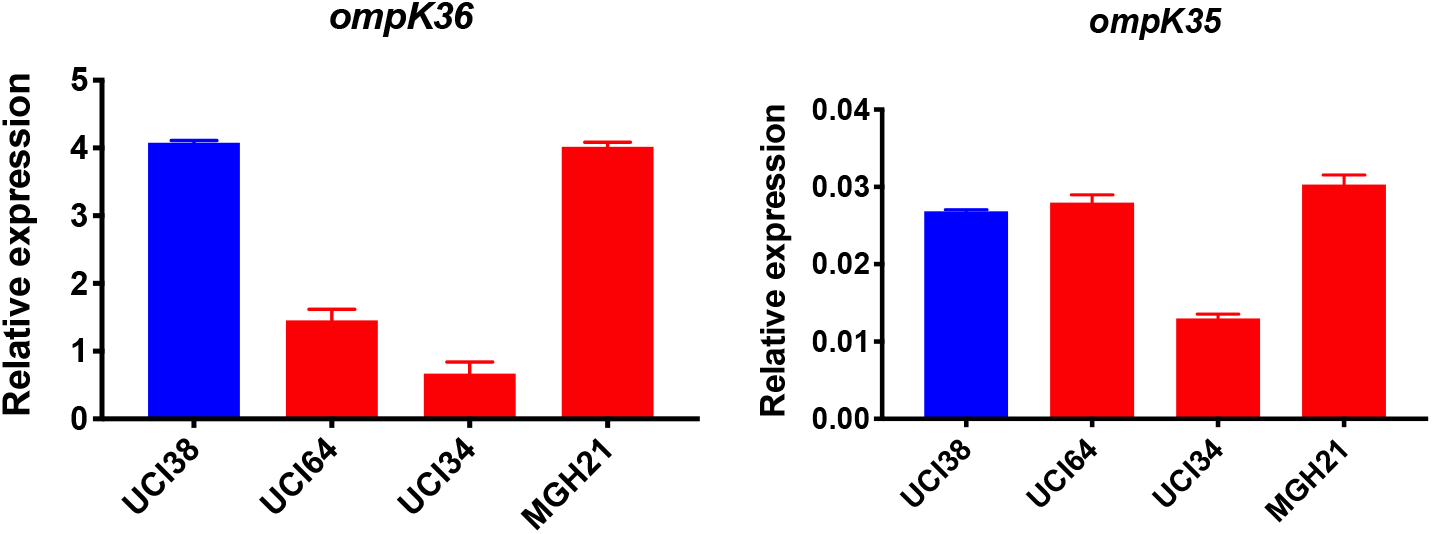
Relative expression of *ompK36* and *ompK35*. RT-qPCR data was normalized to 16S rRNA. Experiments were repeated 3 times and error bars were plotted as standard deviation.

**Supplementary Fig. 5.**
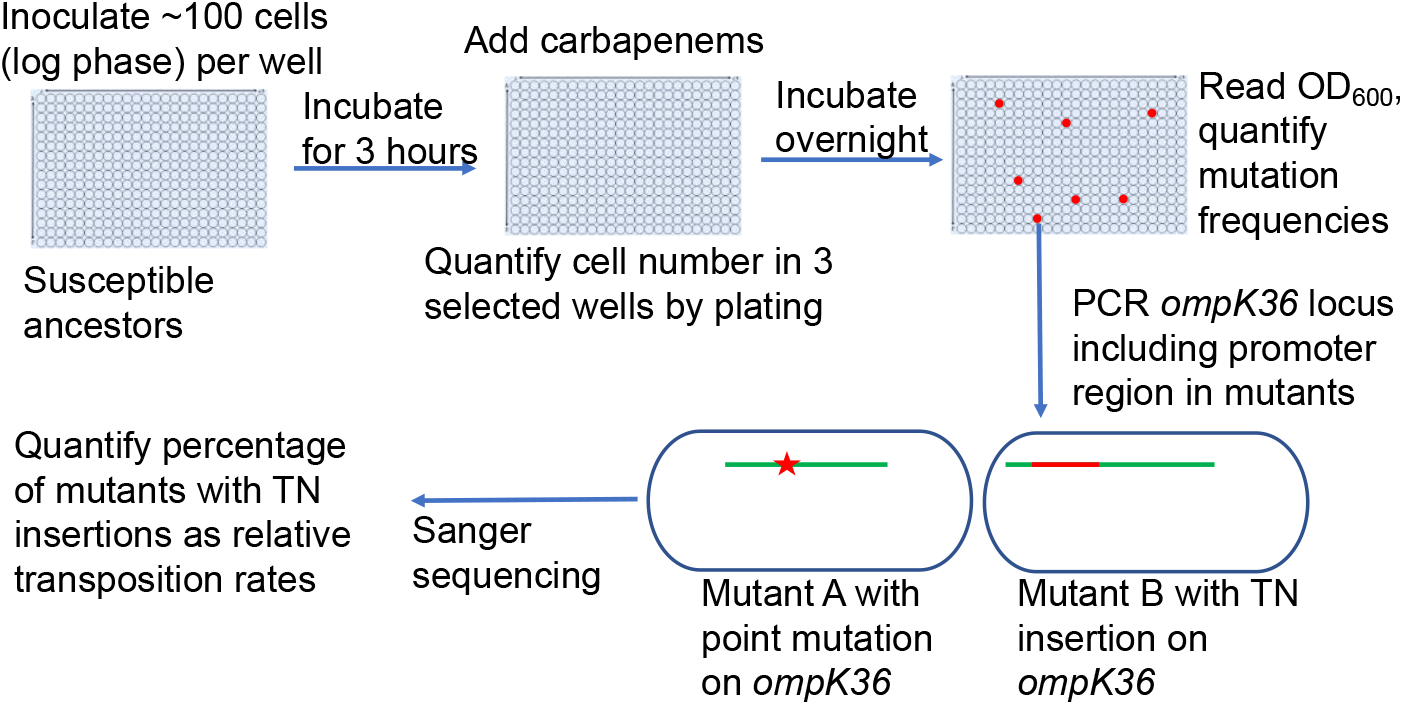
Scheme of the assay for quantification of transposon insertions and SNPs in *ompK36*. Following the isolation of resistant mutants from the modified Luria-Delbrück system, PCR targeting *ompK36* locus, including the upstream 500 bp region, was performed and the PCR products were Sanger sequenced to determine if there were mutations in the targeted region and the types of mutations. Numbers of mutants carrying transposon insertions or SNPs in *ompK36* locus and promoter regions were recorded, and the percentages of mutants with TN insertions or SNP/short INDEL were calculated.

**Supplementary Fig. 6.**
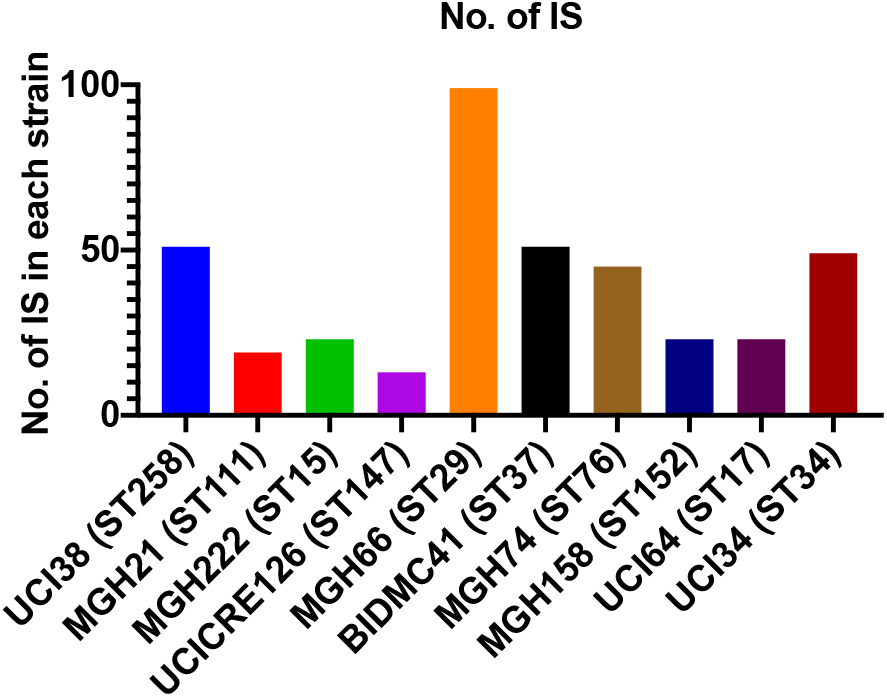
Copy number of ISs in each strain. There is no correlation between the copy number of ISs and the relative level of transposon insertion in *ompK36* locus.

**Supplementary Fig. 7.**
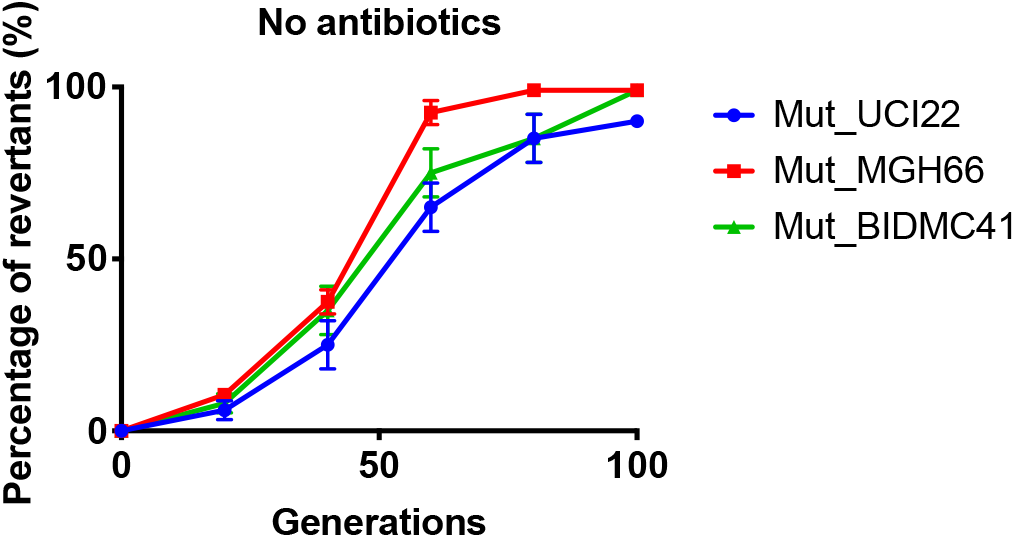
Reversion of TN-insertion mutants derived from UCI22, MGH66 and BIDMC41. In the absence of antibiotics, mutants carrying transposon insertion in the *ompK36* locus and derived from UCI22, MGH66 and BIDMC41 could loss the transposon insertion over ~100 generations. Experiments were repeated 3 times and error bars were plotted as standard deviation.

**Supplementary Fig. 8.**
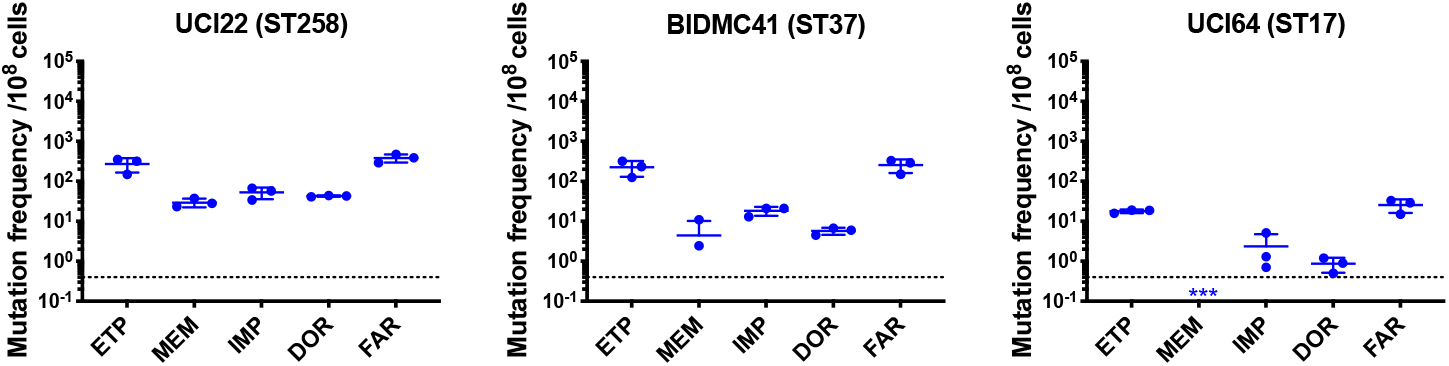
Mutation frequencies of three other isolates under separate treatment with ertapenem (ETP), meropenem (MEM), imipenem (IMP), doripenem (DOR), or faropenem (FAR). Experiments were repeated 3 times and error bars were plotted as standard deviation.

**Supplementary Fig. 9.**
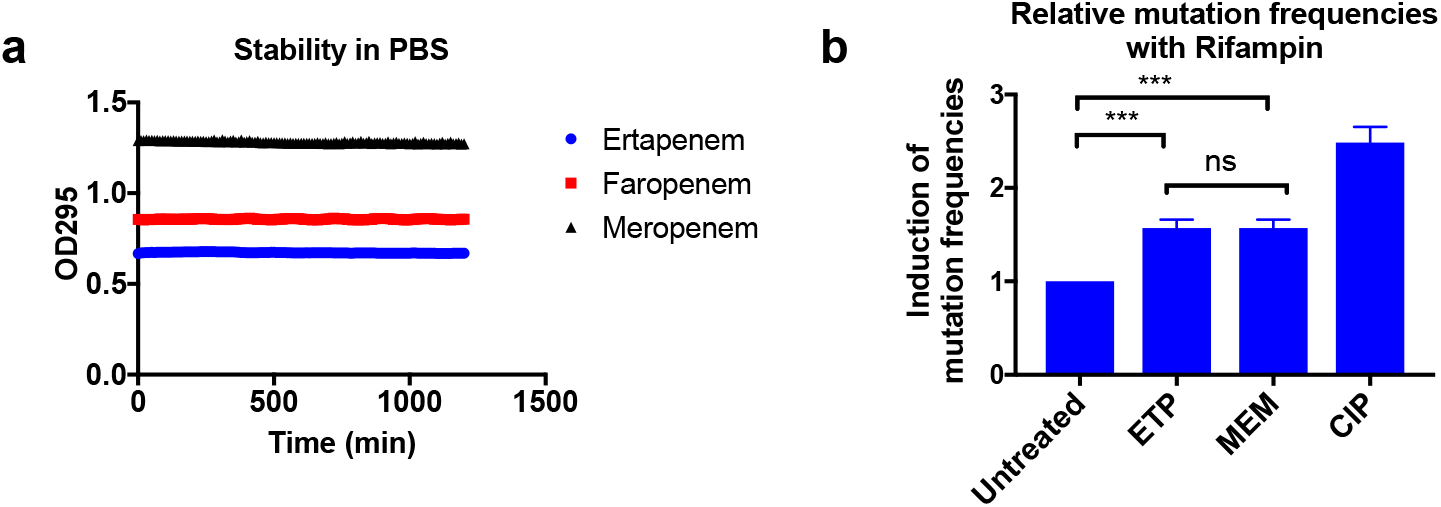
Higher mutation frequencies associated with ertapenem were not due to stability of these drugs or to the induction of mutagenesis. **a**, Stability of ertapenem, faropenem and meropenem in phosphate buffered saline (PBS). Antibiotics were diluted to 0.5 mM in PBS and 100 μl of each antibiotic was used for the assay. OD_295_ was measured every 10 minutes for 20 hours. These three antibiotics are stable for at least 20 hours in our assay condition. **b**, Induction of mutation frequencies under treatment with rifampicin. Bacterial cultures of UCI38 in 384-well plates were pre-treated with ertapenem (ETP), meropenem (MEM) or ciprofloxacin (CIP) at 0.1 x MICs of each drug for 2 hours, then mutation frequencies with rifampicin treatment (50 μg/ml) were measured using the modified Luria-Delbrück system. Ertapenem and meropenem induced mutagenesis to the same degree. Data is plotted as the average of three experiments. Error bars are plotted as the standard deviation. Student t-test was used for statistical analysis to compare the untreated cultures with cultures treated with ertapenem or meropenem.

**Supplementary Fig. 10.**
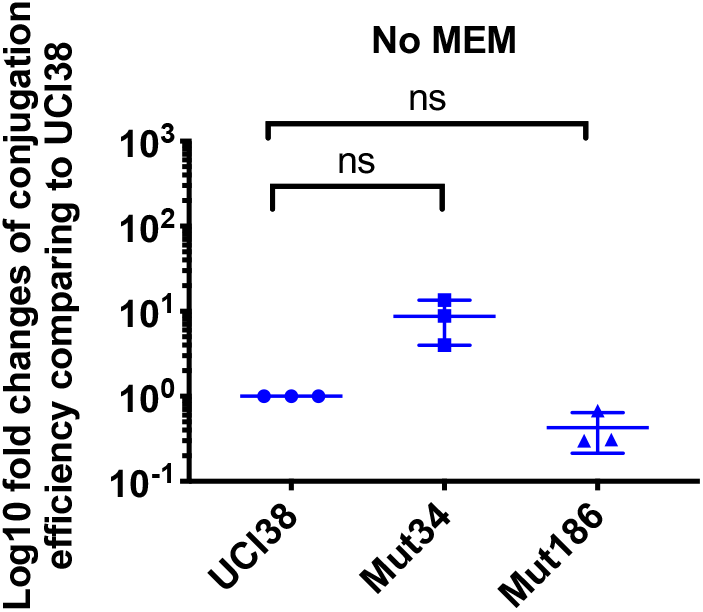
Conjugation efficiencies of UCI38, Mut34 and Mut186 (derived from Mut101 with meropenem treatment) with a *K. pneumoniae* clinical isolate BIDMC45 carrying *bla*_KPC-2_. The conjugation process was conducted in the absence of meropenem. No significant difference was observed in the absence of meropenem between UCI38 and Mut34 (p = 0.11) or between UCI38 and Mut186 (p = 0.16). All experiments were repeated three time. Student t-test was used for statistical analysis to compare between the mutant and the ancestor strain UCI38. Error bars are plotted as standard deviation.

**Supplementary Fig. 11.**
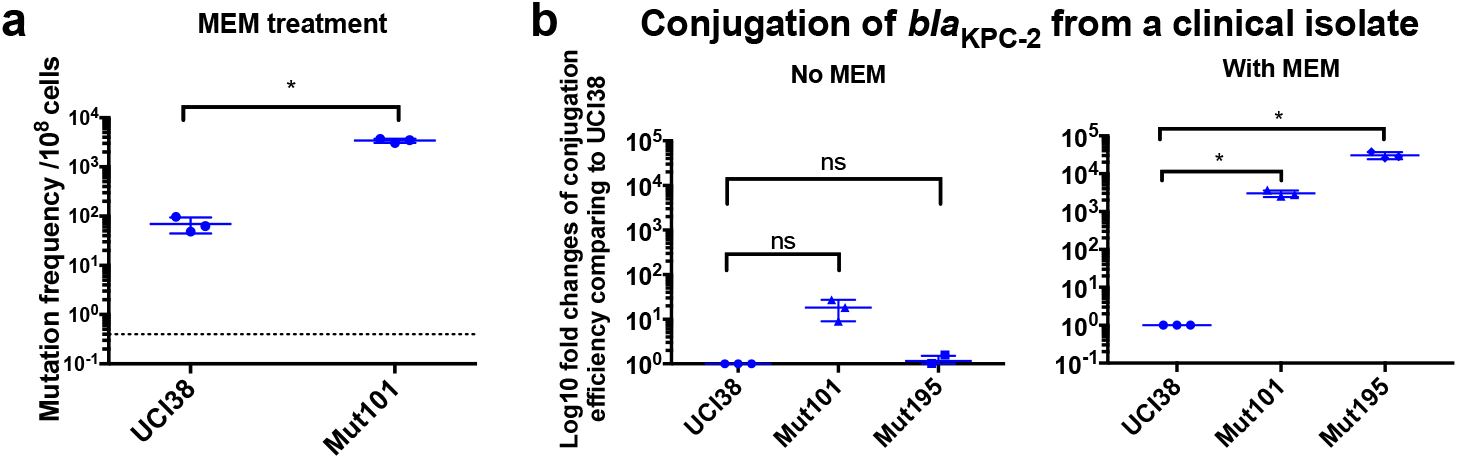
Prior exposure to faropenem promotes the evolution of meropenem resistance. **a**, Mutation frequencies of UCI38 and Mut101 under treatments with meropenem at the concentration of 1.1 x MIC (0.067 μg/ml). Mut101 is a mutant of UCI38 derived from faropenem treatment with the same MIC of meropenem as UCI38 and increased MIC of ertapenem. Mut101 showed significantly higher mutation frequencies than these of the UCI38 with meropenem treatment. **b**, Conjugation efficiencies of UCI38, Mut101and Mut195 (derived from Mut101 with meropenem treatment) with a *K. pneumoniae* clinical isolate BIDMC45 carrying *bla*_KPC-2_. The conjugation process was conducted in the absence or presence of meropenem (0.003 μg/ml). In the presence of meropenem, Mut101 and Mut196 showed higher conjugation efficiencies than these of UCI38. No significant difference was observed in the absence of meropenem between UCI38 and Mut101 (p = 0.08) or between UCI38 and Mut195 (p = 0.5). All experiments were repeated three time. Student t-test was used for statistical analysis to compare the mutation frequencies (**a**) or conjugation efficiencies (**b**) between the mutant and the ancestor strain UCI38. Error bars are plotted as standard deviation.

**Supplementary Fig. 12.**
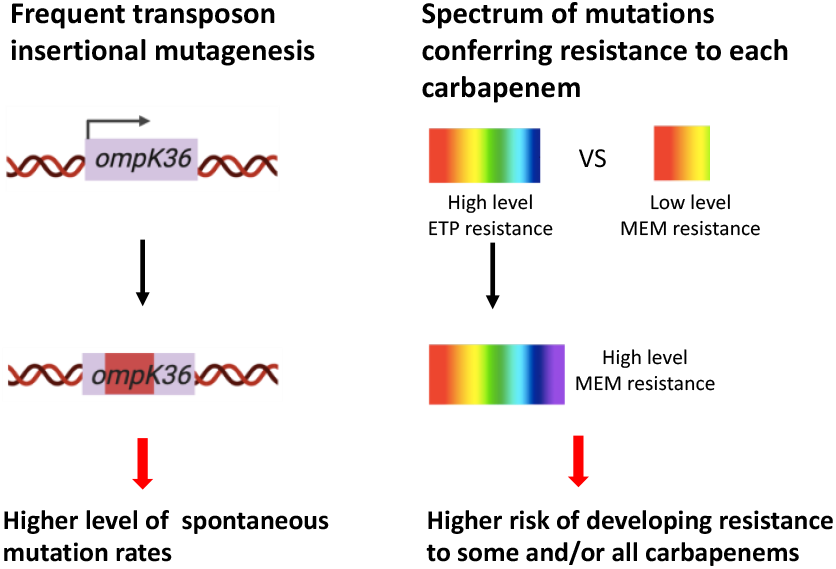
Two genetic determinants of the evolution of carbapenem resistance were identified from this study. On the one hand, high-level transposon insertional mutagenesis facilitates the inactivation of porin genes. On the other hand, a broader spectrum of genetic mutation conferring resistance to ertapenem leads to higher rates of developing resistance with ertapenem treatment; these ertapenem-restricted resistance mutations can serve as stepping-stones to facilitate the development of high-level resistance to all carbapenems.

